# Dysferlin Regulates Cardiac T-tubule Structure and Excitation-contraction Coupling in Isolated Cardiac Myocytes at Rest and in Response to Acute Hypo-osmotic Stress and is Protective Against Arrhythmias in Langendorff-perfused Hearts

**DOI:** 10.1101/2025.08.20.671306

**Authors:** C. J. Quinn, C. J. Booth, K. Smith, E. A. Hayter, E. J. Cartwright, A. W. Trafford, K. M. Dibb

## Abstract

Dysferlin is a membrane-associated protein that supports skeletal muscle function such that mutations in the *DYSF* gene can cause muscular dystrophy. Growing evidence suggests dysferlin regulates cardiac function, but this is less well understood. Tight regulation of the cardiac transverse-(T)-tubule network and excitation-contraction (EC) coupling mechanism is essential for healthy cardiac physiology. Remodelling of T-tubules and the EC coupling mechanism is observed during periods of cardiac stress and pathologies, promoting arrhythmias. However, little is known about how these processes are regulated and any protective mechanisms which may limit detrimental effects. Using a global dysferlin knockout (KO) mouse we have shown that the loss of dysferlin leads to a decrease in T-tubule density and the amplitude and rate of decay of the systolic Ca^2+^ transient but a narrowing of the dyadic cleft. Electrical mapping of *ex vivo* DYSF KO hearts shows they are more susceptible to ventricular arrhythmias. To induce stress, we used hypo-osmotic shock injury (OSI) to damage T-tubule networks in cardiac myocytes, *in vitro*. OSI increased T-tubule fragmentation and caused dysregulation of intracellular Ca^2+^ handling in dysferlin KO cells relative to WT controls. Finally, we observed that a natural decline in WT cardiac dysferlin abundance, which may contribute to the natural age-dependent maladaptive T-tubule remodelling that occurs in the mammalian ventricle. In summary, these findings demonstrate an essential role for dysferlin in cardiac physiology, especially during conditions of stress, which is decreased in normal ageing.

## Introduction

Dysferlin, a 237KDa membrane-associated protein is a major regulator of the skeletal muscle sarcolemma, transverse (T)-tubule network and the excitation-contraction (EC) coupling pathway (Kerr et al. 2013; Lukyanenko et al. 2017; Lukyanenko et al. 2022; Muriel et al. 2022; Quinn et al. 2024). Mutations in the *DYSF* gene cause a specific type of limb-girdle muscular dystrophy known as dysferlinopathy and mounting evidence suggests that dysferlin is also important in the heart (Han *et al*., 2007; Wenzel *et al*., 2007; Chase *et al*., 2009; Wei *et al*., 2015; Hofhuis *et al*., 2020 Paulke et al 2024), which is unsurprising given the extensive structural and functional similarities between skeletal and cardiac muscle.

In cardiac myocytes and skeletal muscle cells, T-tubules are continuations of the sarcolemma that invaginate into the body of the cell to facilitate EC coupling. They do this by allowing the close apposition of the L-type Ca^2+^ channel (LTCC) at the T-tubule membrane with the ryanodine receptor (RyR1/2) on the sarcoplasmic reticulum (SR) Ca^2+^ store. Therefore, the T-tubule network is important for tightly regulated Ca^2+^ release in both tissues (Al-Qusairi and Laporte 2011; Dibb et al. 2022). Alterations in the cardiac T-tubule network and EC coupling mechanism are associated with diseases such as heart failure (HF) and arrhythmias such as atrial fibrillation (AF; (Balijepalli et al. 2003; Cannell et al. 2006; Denham et al. 2018; Dibb et al. 2009; Lenaerts et al. 2009). However, precisely how this tight regulation occurs and any protective mechanisms to maintain it, especially during stress, remain poorly understood. Dysferlin localises to the cardiac T-tubule network in humans (Paulke et al. 2024) and rodents (Hofhuis et al. 2020) and has been shown to drive *de novo* tubule growth in non-cardiac cell lines *in vitro* (Hofhuis et al. 2017). Furthermore, maladaptive cardiac T-tubule remodelling is present in 14-month-old dysferlin-KO mice (Hofhuis et al. 2020).

Recent evidence demonstrates that dysferlin regulates EC coupling in skeletal muscle (Ampong et al. 2005; Flix et al. 2013; Lloyd et al. 2023). Skeletal muscle dysferlin appears to be protective against damage-induced pathological Ca^2+^ handling both *in vitro* and *in vivo* (Kerr et al. 2013; Lukyanenko et al. 2017; Lukyanenko et al. 2022). Whilst less is known about the role of dysferlin in cardiac EC coupling, dysferlin is present within the cardiac dyad (Paulke et al. 2024). Some changes in Ca^2+^ handling have been reported in older murine dysferlin KO cardiac myocytes such as a decreased L-type Ca^2+^ current (*I*_Ca,L_) and SR Ca^2+^ content concurrent with a slower Ca^2+^ transient decay and increased SR Ca^2+^ leak (Hofhuis et al. 2020; Wei et al. 2015); however, our understanding about how dysferlin regulates Ca^2+^ handling in the heart is limited.

In this study we aimed to understand the role of dysferlin in maintaining the structure and function of the EC coupling machinery in health and any protective role during acute and chronic stress where we examined dysferlin’s contribution to normal ageing. We used a global DYSF KO mouse model and found maladaptive T-tubule remodelling in the cardiac myocytes of young DYSF KO mice, which correlated with perturbed intracellular Ca^2+^ handling and increased arrhythmia risk in *ex vivo* Landenforff-perfused hearts. Transmission electron microscopy (TEM) revealed that dysferlin modulates the width of the dyadic cleft.

Using a hypo-osmotic shock injury (OSI) protocol *in vitro* to simulate acute stress we show that the loss of dysferlin renders t-tubules more susceptible to damage and results in dysregulation of Ca^2+^ handling compared to WT controls. Finally, our data suggests that a natural age-related decline in cardiac dysferlin expression may be responsible for the age-dependent decline in mouse cardiac T-tubules. These novel findings demonstrate that dysferlin is a crucial protein for the structural and functional regulation of the cardiac excitation-contraction (EC) coupling machinery and whole heart rhythm.

## Methods

### Animals

All procedures involving animals accord to the United Kingdom (Scientific Procedures) Act of 1986 and the University of Manchester Ethical Review Board. Male and female mice were housed and cared for at the University of Manchester Biological Services Facility. Food and water were available *ad libitum* and lighting was set to 12 hours dark / 12 hours light. The global DYSF KO mouse (B6.129-Dysf^tm1kcam^/J) colony was generated from cryo-recovered embryos obtained from Jax Lab (strain #013149). Experiments were performed using cells or tissue from WT or DYSF KO young (3-5 months) or moderately aged (10-11 months) mice.

### Mouse cardiac myocyte cell isolation

Ventricular cardiac myocytes were isolated using a collagenase and protease digestion technique (Briston et al. 2014). In brief, mice were killed by cervical dislocation and after confirmation of death the hearts were rapidly excised. The aorta was cannulated (22-gauge cannula, Fine Science Tools, USA) and mounted on a Langendorff perfusion system. Hearts were perfused with isolation solution (in mmol/L; NaCl, 134; HEPES, 10; Glucose, 11.1, NaH_2_PO_4_, 1.2; MgSO_4_, 1.2 and KCl, 4) at 3 mL min^-1^ at 37°C containing 0.3 mg/ml Collagenase type I (Sigma, C0130) and 0.03 mg/ml Protease type XIV (Sigma, P5147) for 7.5-10 minutes. The hearts were then perfused with taurine solution (in mmol/L; NaCl, 115; HEPES, 10; Glucose, 11.1, NaH_2_PO_4_, 1.2; MgSO_4_, 1.2; KCl, 4; Taurine, 50 and CaCl, 0.1) at 37°C for 20 minutes. The ventricles were cut into small chunks, gently triturated in taurine solution and the cells were filtered using gauze with a 200 µm pore size (BioDesign Inc. USA).

### T-tubule visualization and quantification

Mouse T-tubules in isolated ventricular myocytes were stained using Di-4-ANEPPS (4 μm/L., Invitrogen, UK) and imaged using a Zeiss 7Live confocal microscope (excitation, 488nm wavelength, emission, 505nm long pass) using an XY pixel dimension of 100nm and a Z-step size of 100nm. Images were deconvolved using the microscope specific derived point spread function using Huygens software (Scientific Volume Imaging, Netherlands). After background correction and thresholding, T-tubule characteristics were quantified using the following equations;

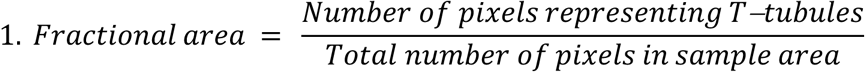

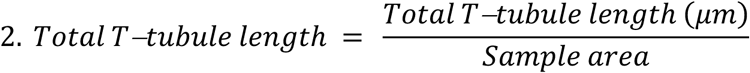

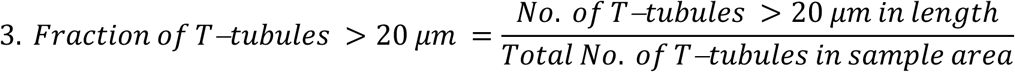

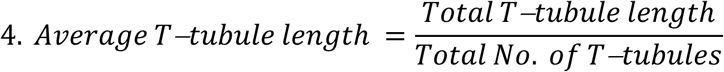

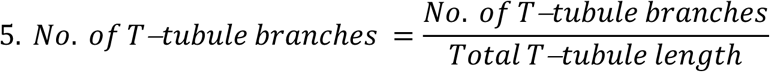

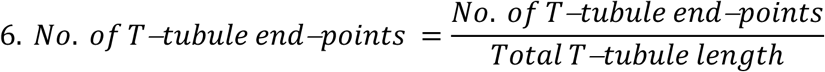

The ratio between transverse and longitudinally oriented tubule branches was calculated using the ImageJ ‘Directionality’ function. Parameters 2-7 were calculated using skeletonised replicas of T-tubule networks.

### Arrhythmia induction via programmed electrical stimulation in Langendorff-perfused hearts

Mice were killed by cervical dislocation and after confirmation of death the hearts were rapidly excised. The aorta was cannulated (22-gauge cannula, Fine Science Tools, USA) and quickly flushed with heparin (1:10) in Krebs-Henseleit (KH) solution containing (in mmol/L; NaCl, 119; NaHCO_3_, 25; KCl, 4; KH_2_PO_4,_ 1.2; MgCl_2_6H_2_O, 1; CaCl_2_2H_2_O, 1.8 and D-glucose, 10). Hearts were mounted on a Langendorff perfusion system and perfused with KH solution at 3 mL min^-1^ at 37 °C. After a 10-minute equilibration period the hearts were subjected to programmed electrical stimulation from the apical region. The hearts were stimulated using an S1-S10 pacing programme where the S1 train consisting of 20 x stimuli (2ms pulse width; 1.5mV) at a basic cycle length (BCL) of 100ms proceeded by an S10 train consisting of 10 x stimuli at a BCL of between 60-20ms decreasing by 10ms per pacing cycle over a total of 5 consecutive cycles. The hearts were then perfused with 100nM isoprenaline in KH solution and after an additional 10-minute equilibration period the S1S10 electrical stimulation programme was repeated. Heart rhythm was monitored using pseudo-ECG (sampling rate = 10KHz) and arrhythmic events were examined in the period immediately after the final S10 stimulus of each of the 5 cycles. The average sustained only, triggered only and total arrhythmia scores was then calculated for each heart according to a modified Lambeth scoring method (Walker et al. 1988), where the score increases with in line with the severity of the arrhythmic event as follows; single PVC = 1, two PVCs = 3, three PVCs = 4, VT (four or more PVCs) < 1s = 5 and VT > 1s = 6.

### Osmotic shock injury assay

Myocytes were subjected to *in vitro* hypo-osmotic shock injury as per Moench et al., 2013. Cells were bathed in a normo-osmotic Tyrode solution (1.0 Tyr) containing (in mmol/L; NaCl, 140; HEPES, 8.6; Glucose, 10; KCl, 4; MgCl, 1 and CaCl 0.3) for 10 minutes. The cells were then submerged in a hypo-osmotic Tyrode solution (0.7 Tyr) containing 0.7x the NaCl concentration of the 1.0 Tyr solution (98 mM) for 7 minutes. For live Ca^2+^ imaging experiments the cells were returned to 1.0 Tyr solution and imaged immediately at 37°C. For T-tubule analysis, the cells were treated with Di-4-ANEPPS as described previously (Dibb et al. 2009) after returning to 1.0 Tyr solution and imaging commenced after 10 minutes.

### Protein quantification

Protein abundance was examined using established Western blot techniques (Towbin et al. 1979). In brief, protein samples were treated with an anti-dysferlin primary (1°) antibody (B13106; LS Bio; 1:100) overnight at 4°C and a secondary (2°) antibody (LiCOR anti-rabbit W800C) for 1 hour at RT. Stained membranes were imaged using a LI-COR Odyssey CLx imaging system. Dysferlin protein abundance was normalised to a LI-COR total protein stain for each gel well.

### Intracellular Ca^2+^ measurements

#### Basic intracellular Ca^2+^ handling parameters

Cardiac myocytes were loaded with Fura-2 AM (Life Technologies, USA; 1 µM) ratiometric Ca^2+^ indicator. Cells were perfused with Normal Tyrode’s solution containing (in mmol/L); NaCl, 140; HEPES, 8.6; Glucose, 10; KCl, 4; MgCl, 1; and CaCl 1) at 37°C. Field stimulation was used to excite cells according to a defined pacing sequence (0.5, 1, 0.5, 2, 0.5 Hz) and the Fura-2 signal was recorded continuously. Fluorescence was excited at 340 and 380nm and emitted fluorescence was detected at 510 ± 10 nm. After subtracting the background fluorescence, the ratio of light excited at 340 nm to that excited at 380nm was used to measure changes in intracellular Ca^2+^. Pacing at each rate was continued until steady-state was reached and parameters of the Ca^2+^ transient were calculated from a 10-beat average. For 0.5 Hz, mean values from each of the three 0.5Hz pacing cycles in the same cell were used. The rate constant of decay (1/tau) of the Ca^2+^ transient (*k*_sys_) was calculated in Clampfit V11.0.3 by fitting a standard single exponential curve. 10mM caffeine was rapidly applied to measure sarcoplasmic reticulum Ca^2+^ content (Hutchings et al. 2022). Na^+^/Ca^2+^ exchanger (NCX) activity was approximated by measuring the rate constant of decay of a single exponential fit to the 10 mM caffeine-induced Ca^2+^ transient *(k*_caff_). The contribution of SERCA to the rate constant of decay of the Ca^2+^ transient (*k*_sr_) was calculated using the following equation;

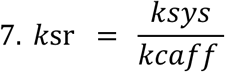

Finally, fractional SR Ca^2+^ release at each pacing rate was calculated using the following equation;

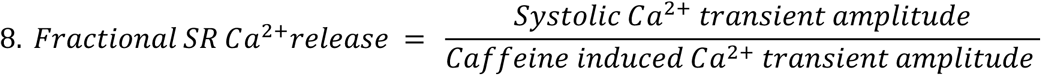

### Ca^2+^ spark frequency, Ca^2+^ waves / ectopic beats & dyssynchrony of Ca^2+^ release

Ventricular cardiac myocytes were loaded with Fluo-5F (5 µM). Cells were stimulated at room temperature using field electrodes at 0.5 Hz for 10 seconds followed by a 10 second unstimulated rest period whilst Ca^2+^ was measured continuously in Normal Tyrode’s solution with or without 100 nM isoprenaline. To limit bleaching of the Ca^2+^ indicator, one-dimensional-line scan was performed with a Zeiss 7Live microscope 488nm wavelength and 505 long pass emission settings at 1 KHz. Ca^2+^ spark frequency was analysed using custom analysis software (MatLab). Following background subtraction Ca^2+^ transient amplitude was calculated in ImageJ as the ratio of the peak fluorescence to baseline fluorescence of each stimulation using normalized F/F0 line scans. The Ca^2+^ dyssynchrony index was calculated using custom code (Matlab).

### Electron Microscopy

To visualise subcellular structures in ventricular myocytes with nanometre resolution we used standard transmission electron microscopy (Winey et al. 2014). Small chunks (0.5mm^3^) of mouse left ventricular myocardium were prepared as described previously (Caldwell et al. 2024). For TEM image acquisition, the samples were removed from the moulds and cut into ultrathin sections (60-80 nm) before being placed onto mesh grids. Images were then taken at 28000x magnification using a TALOS L120C TEM with a Ceta camera. TEM images were taken from a fraction of 2-3 randomly selected cells per animal.

The InteredgeDistance_V1.0.1 ImageJ macro was used to accurately measure the dyadic cleft width. The dyadic portion of the tubule and SR membranes were delineated and the InteredgeDistance_V1.0.1 macro made 50 individual measurements across the length of the dyadic cleft. From these measurements, the average and minimum dyadic cleft widths and standard deviation (SD) were obtained for each dyad. The coefficient of variation was used to examine interdyadic width variability. Due to variation in the orientation of the ultrathin slices the narrowest luminal T-tubule width was used to determine t-tubule diameter. T-tubule to Z-line distance was measured manually using ImageJ as described previously by Zhang et al., 2013. The T-tubule coupling ratio was calculated using the following equation;

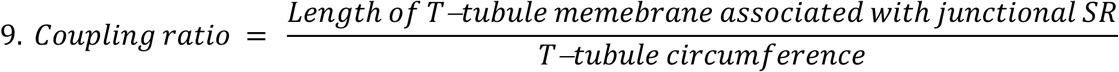

### Statistical Analysis

Data are presented as mean ± standard error of the mean (SEM) for *n* observations/*N* experiments (animals). Where multiple observations (*n*; such as multiple individual cell measurements from the same heart) were made from the same animal (*N*) linear mixed modelling (SPSS Statistics. IBM, USA) was used to account for the nested (clustered) experimental design (Caldwell et al. 2014). In instances where the data were not normally distributed (unless normal distribution could not be achieved), data were transformed using Log10 or reciprocal (1/x) to achieve normal distribution prior to linear mixed modelling. Statistical significance was also determined using the following tests (as detailed in the corresponding figure legends); One-way ANOVA with either a; Tukey’s multiple comparisons or a Fisher’s least significant difference (LSD) post hoc test, Fisher’s Exact test, two-way ANOVA, Mann-Whitney test, paired two-tailed t-test, unpaired two-tailed t-test or a propagation of errors calculation using a one-tailed t-test. *P* values of <0.05 show statistical significance and exact *P* – values are stated if *P =* > 0.0001.

## Results

### Dysferlin regulates cardiac T-tubule structure

To investigate the role of dysferlin in the regulation of mammalian cardiac T-tubules, we used confocal imaging to visualise the T-tubule network in ventricular myocytes isolated from 3-5-month-old WT and DYSF KO mice (**Fig. 1A & B**). Multiple T-tubule characteristics were calculated using ImageJ from skeletonized T-tubule networks where the sarcolemma had been digitally removed to focus on T-tubules only (**Fig. 1Ai & Bi**). T-tubule density was decreased in DYSF KO cells compared to WT controls both by measuring the T-tubule fractional area (**Fig. 1C**) and the total T-tubule length (**Fig. 1D**). There was no difference in T-tubule orientation between WT and DYSF KO cells (**Fig. 1E**). T-tubules were more fragmented in DYSF KO cells compared to WT cells as indicated by an increase in the number of individual T-tubule structures (**Fig. 1F**), a decrease in the average T-tubule length (**Fig. 1G**) and a decrease in the occurrence of large, interconnected T-tubules (**Fig. 1H**). A trend towards a decrease in the number of T-tubule branches (**Fig. 1J**), and a reduction in the number of interconnecting T-tubule junctions (**Fig. 1K**) may account for the increased fragmentation observed in the DYSF KO T-tubule network. To examine the ultrastructure of the T-tubule network we used transmission electron microscopy. At the region of the Z-line, all the WT T-tubules we identified appeared ordered with a healthy morphology (**Fig. 1L**), whereas we observed examples of T-tubule disorder in 25% (2/8) of DYSF KO myocardial tissue samples (**Fig. 1M & N**). Given the less dense and more fragmented nature of the DYSF KO T-tubules, we next sought to establish if the association between the T-tubules and the SR was altered.

**Figure 1.**
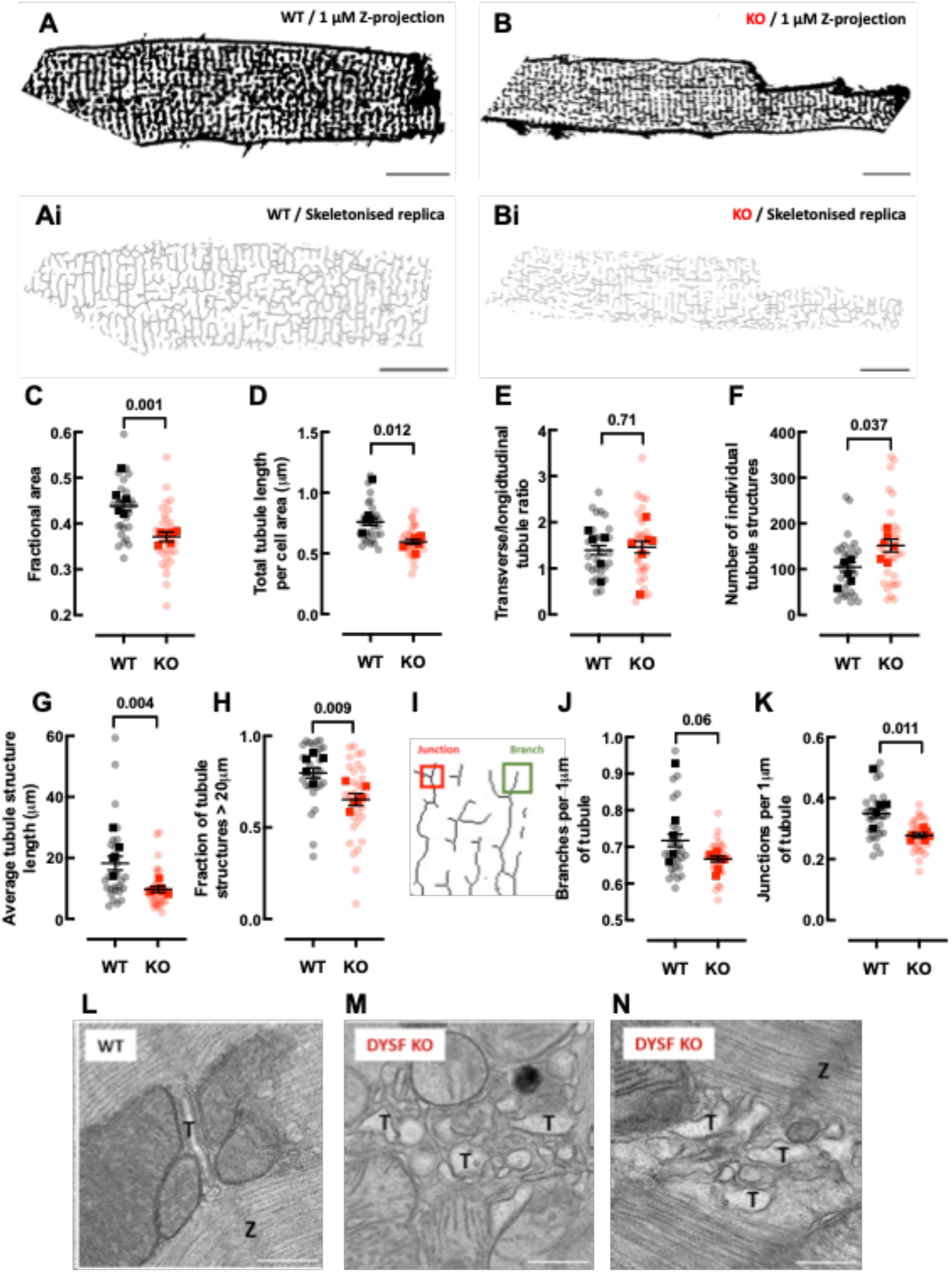
Dysferlin loss decreases T-tubule density, fragmenting and decreasing connectivity in remaining tubules. **A & B**, Representative composite confocal images of the sarcolemma and the T-tubule network stained with Di-4-ANEPPS in isolated WT and DYSF KO ventricular myocytes. **Ai** & **Bi**, Skeletonised replicas of the T-tubule networks shown in **A** & **B**, respectively. **A-Bi**, Scale bars = 15µm. **C**, T-tubule density measured by fractional area. **D**, Total tubule length normalized to cell area (µm). **E**, Transverse/longitudinal tubule ratio. **F**, Number of individual tubule structures. **G**, Average tubule structure length (µm). **H**, Fraction of tubule structures over 20 µm in length. **I**, Zoom panel showing T-tubules outlined in **Ai** demonstrating junctional and branch structures. **J**, Number of T-tubule branches per 1 µm of tubule. **K**, Number of T-tubule junctions per 1 µm of tubule. *N* = 5-6 mice (opaque data points), *n* = 27-29 cells (translucent data points). Linear mixed model analysis was used to determine statistical significance. Data is presented as mean ± SEM. Example electron micrographs at 28000x magnification of T-tubule cross sections and T-tubule disorder in WT (**L**) and DYSF KO mouse left ventricular myocardial tissue (**M** & **N**). Scale bars = 200nm. T = suspected T-tubule lumen. Z = Z-disc.

### The width of the dyadic cleft is narrowed in DYSF KO myocytes

The limited resolution of confocal imaging precludes its use as a method for studying the ultrastructure of T-tubules or the cardiac dyad. Therefore, we used 2D transmission electron microscopy (**Fig. 2A & B**). The loss of dysferlin did not affect T-tubule diameter (**Fig. 2C**) or the positioning of the T-tubules at the Z-line (**Fig. 2D**). Similarly, the fraction of the T-tubule membrane associated with junctional SR was not different between WT and DYSF KO myocytes (**Fig. 2E**). We observed a decrease in the width of the dyadic cleft in DYSF KO cells compared to WT controls (**Fig. 2F & G**) without any change in the intra- or inter-dyadic cleft width variability (**Fig. 2H & I**). These findings suggest a uniform narrowing of the dyadic cleft in DYSF KO cells, which to the best of our knowledge, is the first ever report of a narrowing of the cardiac dyad. Given the loss of T-tubule density but narrowing of the dyad, we then set out to determine if the loss of dysferlin altered intracellular Ca^2+^ handling.

**Figure 2.**
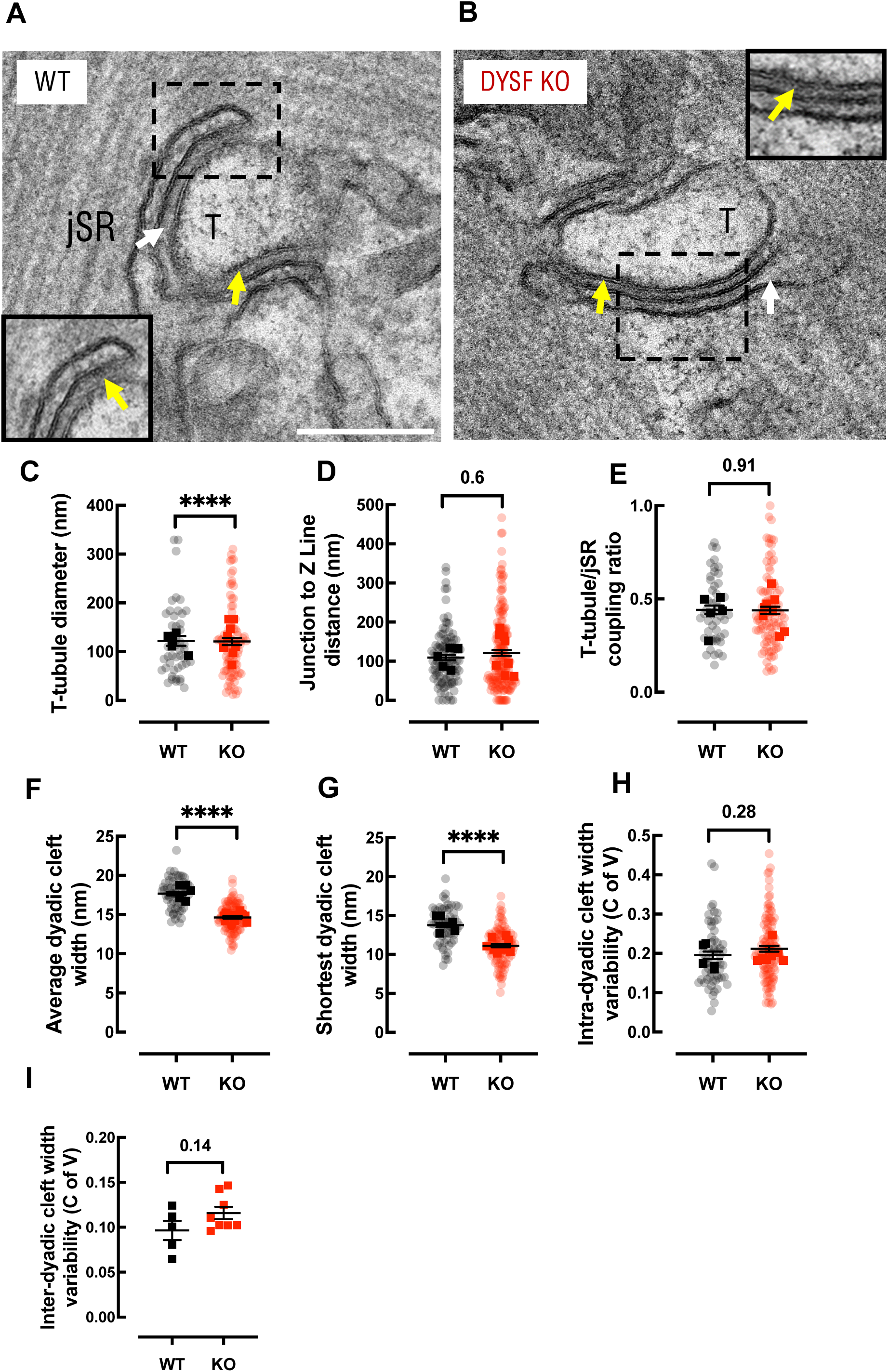
Dysferlin regulates the width of the dyadic cleft. **A & B**, Representative transmission electron micrographs showing a cross section of a mouse left ventricular T-tubule (T) forming a dyadic cleft (yellow arrows) with the junctional sarcoplasmic reticulum (jSR; white arrows) in ultrathin myocardial sections from a WT and DYSF KO mouse, respectively. **A & B** Scale bars = 200 nm. **C**, T-tubule luminal diameter (nm). **D**, Junction to Z-line distance (nm). **E**, T-tubule/jSR coupling ratio. **F**, Average dyadic cleft width (nm). **G**, Shortest dyadic cleft width (nm). **H**, Intra-dyadic cleft width variability measured by the coefficient of variation (CofV). *N* = 5-8 mice (opaque data points), *n* = 46-174 measurements (translucent data points). Linear mixed model analysis was used to determine statistical significance. **I**, Inter-dyadic cleft variability (C of V). *N* = 5-8 mice. Statistical significance was determined using a nested t-test. Data is presented as mean ± SEM. **** - *P* = <0.0001.

### Dysferlin KO leads to altered intracellular Ca^2+^ handling

Epifluorescence imaging with field stimulation was used to examine Ca^2+^ handling. Representative stimulated Ca^2+^ transients and superimposed mean Ca^2+^ transients in response to 0.5, 1 and 2 Hz stimulation are shown in **Figures 3A & B**. As expected, increasing stimulation frequency resulted in an increase in diastolic Ca^2+^ in both WT and DYSF KO cells but diastolic Ca^2+^ was comparable between genotypes at all stimulation rates (**Fig. 3C**). There was an overall decrease in Ca^2+^ transient amplitude in DYSF KO relative to WT cells and pairwise comparisons also showed a trend towards a decrease in Ca^2+^ transient amplitude at higher rates of stimulation in DYSF KO cells (**Fig. 3D**). This was accompanied by a decrease in the rate of decay of the systolic Ca^2+^ transient (*k*_sys_) at all rates in DYSF KO cells (**Fig. 3E**). Representative superimposed caffeine-induced Ca^2+^ transients are shown in **Figure 3F**. Whilst the amplitude of the caffeine-induced Ca^2+^ transient was unchanged with genotypes (**Fig. 3G**), there was an overall decrease in the fractional SR Ca^2+^ release in DYSF KO cells compared to WT cells (**Fig. 3H**). The rate of decay of the caffeine-induced Ca^2+^ transient (*K*_caff_) was decreased in DYSF KO cells at all stimulation rates (**Fig. 3I**). Calculation of the contribution of SERCA to systolic Ca^2+^ transient decay (*K*_sr_), revealed *K*_sr_ was also decreased in DYSF KO cells at all stimulation rates (**Fig. 3J**). Therefore, the decrease in *K*_sys_ was due to a decrease in both *K*_caff_ and *K*_sr_. In summary, the loss of dysferlin expression leads to decreased Ca^2+^ transient amplitude and SR fractional Ca^2+^ release together with impaired removal of cytosolic Ca^2+^ under baseline conditions.

**Figure 3.**
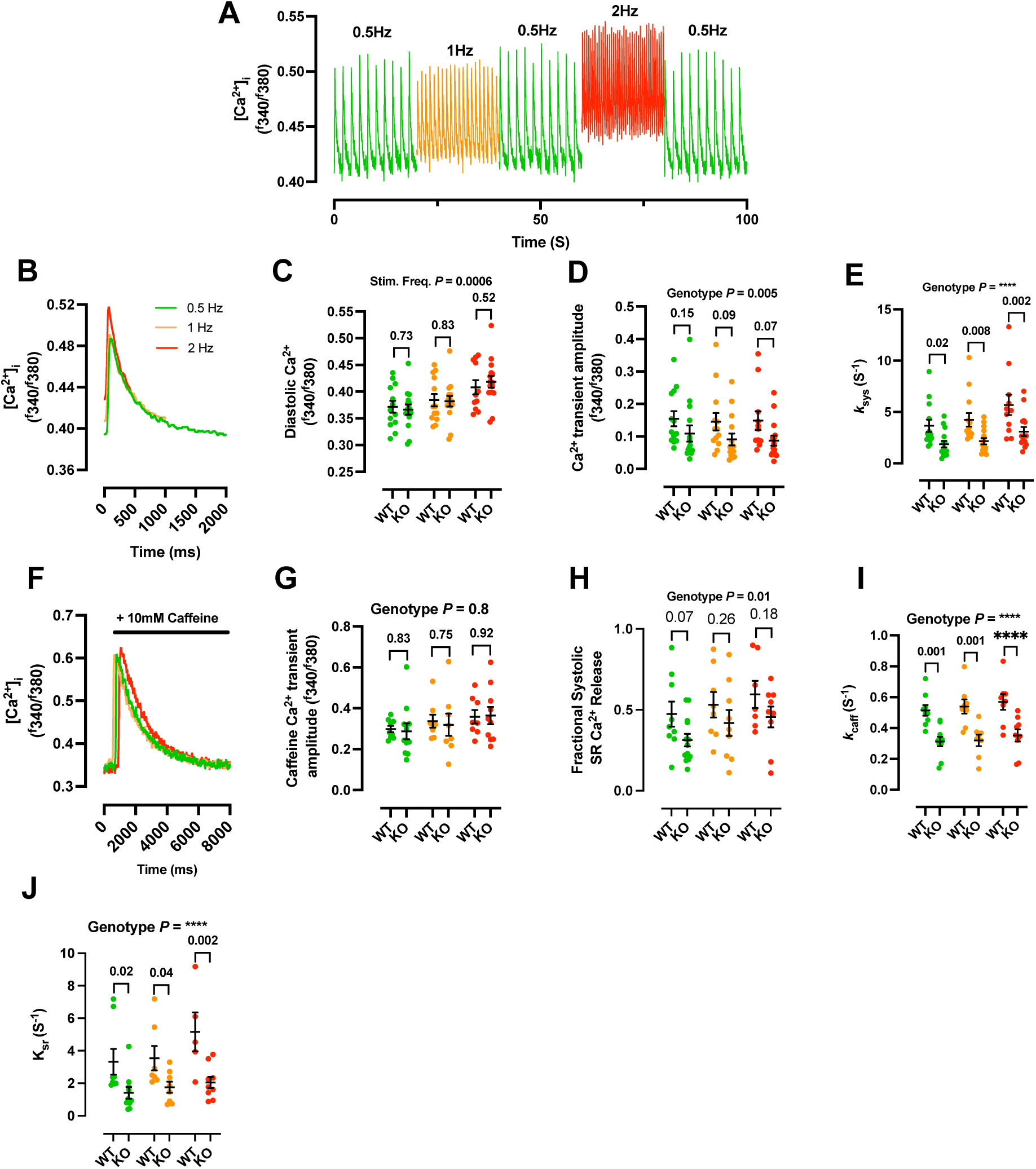
Dysferlin loss causes decreases in Ca^2+^ transient amplitude, fractional SR Ca^2+^ release and SERCA dependent and independent components of the rate of Ca^2+^ removal. Isolated cardiac myocytes were loaded with a Ca^2+^ sensitive dye (Fura-2) and stimulated using field stimulation at 0.5 (green), 1 (orange) and 2 Hz (red) sequentially in 1 mM Ca^2+^ Normal Tyrode’s solution. **A**, Representative stimulated Ca^2+^ transients from a WT cardiac myocyte demonstrating experimental pacing sequence ([Ca^2+^]_I_ (^f^340/^f^380)). **B**, Representative Ca^2+^transients from a WT cell (10-beat average) at each stimulation frequency. **C**, diastolic Ca^2+^ **D,** Ca^2+^ transient amplitude and **E,** Rate constant of decay (1/t; RoD) of the Ca^2+^ transient (K_sys_ S^-1^). **F**, Representative 10 mM caffeine-induced Ca^2+^ transients from a WT cell. **G**, 10 mM caffeine-induced Ca^2+^ transient amplitude **H,** Fractional systolic SR Ca^2+^ release **I**, RoD of the caffeine-induced Ca^2+^ transient (Kcaff S^-1^) and **J**, SERCA-mediated RoD (Ksr S^-1^). **C**, **D**, **E**, **G**, **H**, **I** & **J**, *N* = 4-6, *n* = 5-15. Statistical significance was determined using a one-way ANOVA with a post hoc Fisher’s least significant difference (LSD) test. **C, D, G, J, H, I & J** A two-way ANOVA was used to determine the overall effect of genotype or frequency of stimulation for each parameter. Data is presented as mean ± SEM. **** - *P* = <0.0001.

To determine if the T-tubule disruption upon dysferlin loss decreased the synchronicity of Ca^2+^ release, confocal line scan experiments were performed. Despite a decreased T-tubule density in DYSF KO cells (**Fig. 1**), the synchronicity of the rise in the Ca^2+^ transient was unaltered when stimulated at 0.5Hz (**Supplementary Figure 2**). In the same experiments we examined Ca^2+^ sparks to determine if the decrease in SR fractional release in the DYSF KO was associated with altered RyR Ca^2+^ release. However, we found that Ca^2+^ spark frequency was similar between genotypes (**Supplementary Figure 2**).

### Dysferlin is protective against ventricular arrhythmias in *ex vivo* hearts

We subjected ex vivo Langendorff-perfused hearts to S1S10 programmed electrical stimulation to determine whether the observed maladaptive remodelling of the T-tubule network and EC coupling mechanism in DYSF KO cardiac myocytes was sufficient to alter heart rhythm. Heart rhythm was monitored using pseudo-ECG, and arrhythmic events were scored using a modified Lambeth scoring system (**Fig. 4A-Aiv**). Dysferlin decreased the number of total arrhythmic events in the presence of 100nM isoprenaline using the S1S10 pacing programme (**Fig. 4B** & **C**). To examine this mechanism more closely, we quantified triggered arrhythmic events (1, 2 or 3 PVCs) and sustained arrhythmias (VT; defined as 4 or more PVCs) separately. From this, we determined that the overall increased arrhythmia risk in DYSF KO hearts when exposed to isoprenaline was due to an increased propensity for VT (**Fig. 5D-F**) rather than any change in triggered events, which was comparable in DYSF KO and WT hearts (**Fig. 5G-I**). These findings demonstrate that dysferlin can protect the heart from cardiac arrhythmias. Since arrhythmias often occur during conditions of stress we next set out to determine any protective role of dysferlin during cardiac stress.

**Figure 4.**
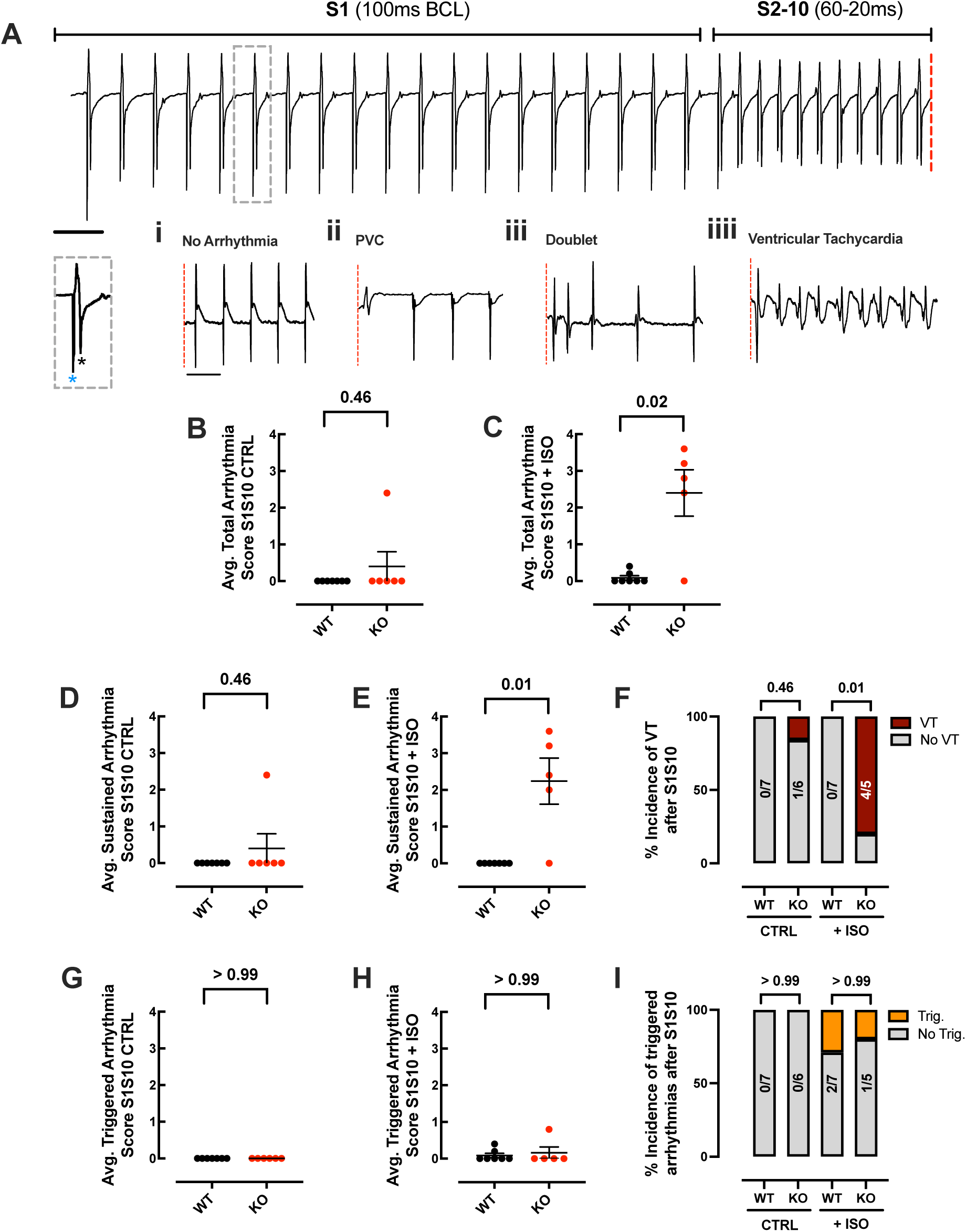
Dysferlin loss leads to an increased risk of arrhythmias in response to programmed electrical stimulation (PES) in Langendorff-perfused hearts. **A**, Example S1-S10 PES protocol where the S1 train = 20 x stimuli at a basic cycle length (BCL) of 100ms is proceeded by an S10 train = 10 x stimuli at a BCL of 60-20ms decreasing by 10ms per pacing cycle over a total of 5 consecutive cycles. Zoom panel illustrates a single S1 stimulation artefact (pulse width = 2ms) indicated by a blue ‘*’ leading to a stimulated ventricular complex indicated by a black ‘*’. Heart rhythm and the manifestation of ventricular arrhythmic events were monitored using pseudo-ECG during the period immediately after the final stimulus of the S10 train (red line). These phenomena include; **i**, a return to sinus rhythm indicating no arrhythmias (score = 0), **ii**, a single premature ventricular complex (PVC; score = 1), **iii**, 2 x consecutive PVCs (Doublet; score = 3) or **iv**, sustained ventricular tachycardia (VT; score = 5 if episode of VT persisted for <1 second and score = 6 if VT episode persisted for >1 second.). **A-Aiv**, Scale bars = 125ms. **B** & **C,** Average total arrhythmia score after S1S10 PES – and + 100nM isoprenaline (ISO), respectively. **D** & **E**, Average arrhythmia score after S1S10 PES for sustained arrhythmias (VT) only, – and + ISO, respectively. **F**, Percentage incidence of VT – and + ISO. **G** & **H,** average arrhythmia score for triggered arrhythmias (PVC or Doublet) only, - and + ISO, respectively. **I**, Percentage incidence of triggered arrhythmias – and + ISO. **B**, **C**, **D**, **E**, **G** & **H**, Statistical significance was determined using a Mann-Whitney test. **F** & **I**, Statistical significance was determined using a Fisher’s Exact test. Data points represent a modified Lambeth arrhythmia score for a single heart. *N* = WT – 7, KO – 5. Data is presented as mean ± SEM.

**Figure 5.**
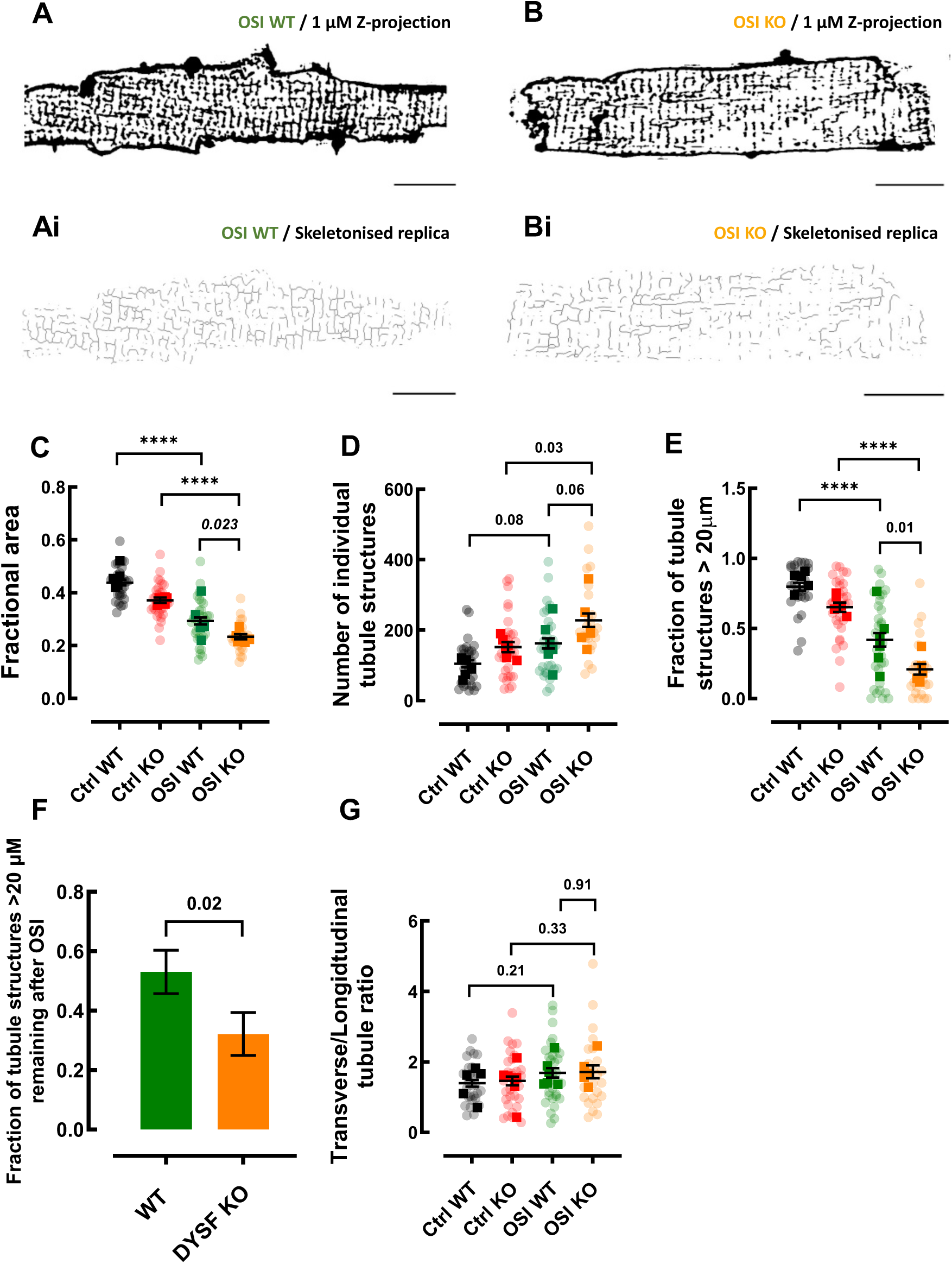
*In vitro* osmotic shock injury causes increased T-tubule fragmentation in DYSF KO cardiac myocytes. **A & B**, Representative composite confocal images of the sarcolemma and the T-tubule network stained with Di-4-ANEPPS in isolated WT and DYSF KO mouse ventricular myocytes following OSI. **Ai & Bi**, Skeletonised replicas of the T-tubule networks shown in **A** & **B**. **A-Bi**, Scale bars = 15 µm. **C**, T-tubule fractional area. **D**, Number of individual tubule structures. **E**, Fraction of tubule structures over 20 µm in length. **F**, Fractional reduction in the number of tubule structures over 20 µm in length after OSI. **G**, Transverse/longitudinal tubule ratio. *N* = 5-6 mice (opaque data points), *n* = 23-30 cells (translucent data points). **C, D, E** & **G,** Linear mixed model analysis was used to determine statistical significance. **F**, Statistical significance was determined using a propagation of errors calculation and a one-tailed t-test. Data is presented as mean ± SEM. **** - *P* = <0.0001.

### DYSF KO T-tubules become more fragmented after osmotic shock injury (OSI)

To understand the relationship between dysferlin and the T-tubule network in response to acute stress, we subjected isolated mouse ventricular myocytes to osmotic shock injury (OSI) *in vitro*, to study the relationship between dysferlin and the T-tubule network in response to acute stress. The T-tubule network was visualised using confocal z-stacks in isolated myocytes from WT and DYSF KO mice after OSI (**Fig. 5A & B**). T-tubule network properties were quantified from skeletonized T-tubule network replicas using ImageJ (**Fig. 5Ai & Bi**) and compared to control data presented in **Figure 1**. OSI caused a similar reduction in overall T-tubule density in both WT and DYSF KO cells; thus, the decreased T-tubule density in DYSF KO cells persisted after OSI (**Fig. 5C**). However, in WT cells OSI had no effect on the number of T-tubule structures, whereas in the DYSF KO, OSI increased the number of tubule structures in DYSF KO cells, consistent with tubule fragmentation (**Fig. 5D**). Consistent with enhanced fragmentation of T-tubules in DYSF KO cells, the decrease in the fraction of large, interconnected T-tubules observed in WT and DYSF KO cells (**Fig. 5E**) was more severe in DYSF KO cells (**Fig. 5F**). We did not observe a change in T-tubule orientation after OSI in either genotype (**Fig. 5G**). These data show that dysferlin acts to protect the mammalian cardiac T-tubule network against acute stress *in vitro*.

### DYSF-KO cardiac myocytes are unable to adapt to changes in extracellular Ca^2+^ concentration and show increased systolic Ca^2+^ release after OSI

Next, to determine if dysferlin protects Ca^2+^ handling during periods of damage, we investigated the impact of OSI on intracellular Ca^2+^ handling in cardiac myocytes using epifluorescence. In contrast to baseline Ca^2+^ handling experiments, which were performed in the standard 1mM Ca^2+^ Tyrode’s solution (**Fig. 3**), OSI Ca^2+^ measurements were performed in 0.5mM Ca^2+^ Tyrode’s to avoid Ca^2+^ overload in low Na^+^ concentration due to reverse mode Na^2+^/Ca^2+^ exchanger (NCX) function during the hypo-osmotic period (Akar et al. 2005). The OSI protocol is shown in **Figure 6A** where ‘*’ indicates the time points at which Ca^2+^ measurements were made in **Figures 6E-G**. Example average Ca^2+^ transients from WT and DYSF KO cells post-OSI are shown in **Figures 6B** & **C**, respectively.

**Figure 6.**
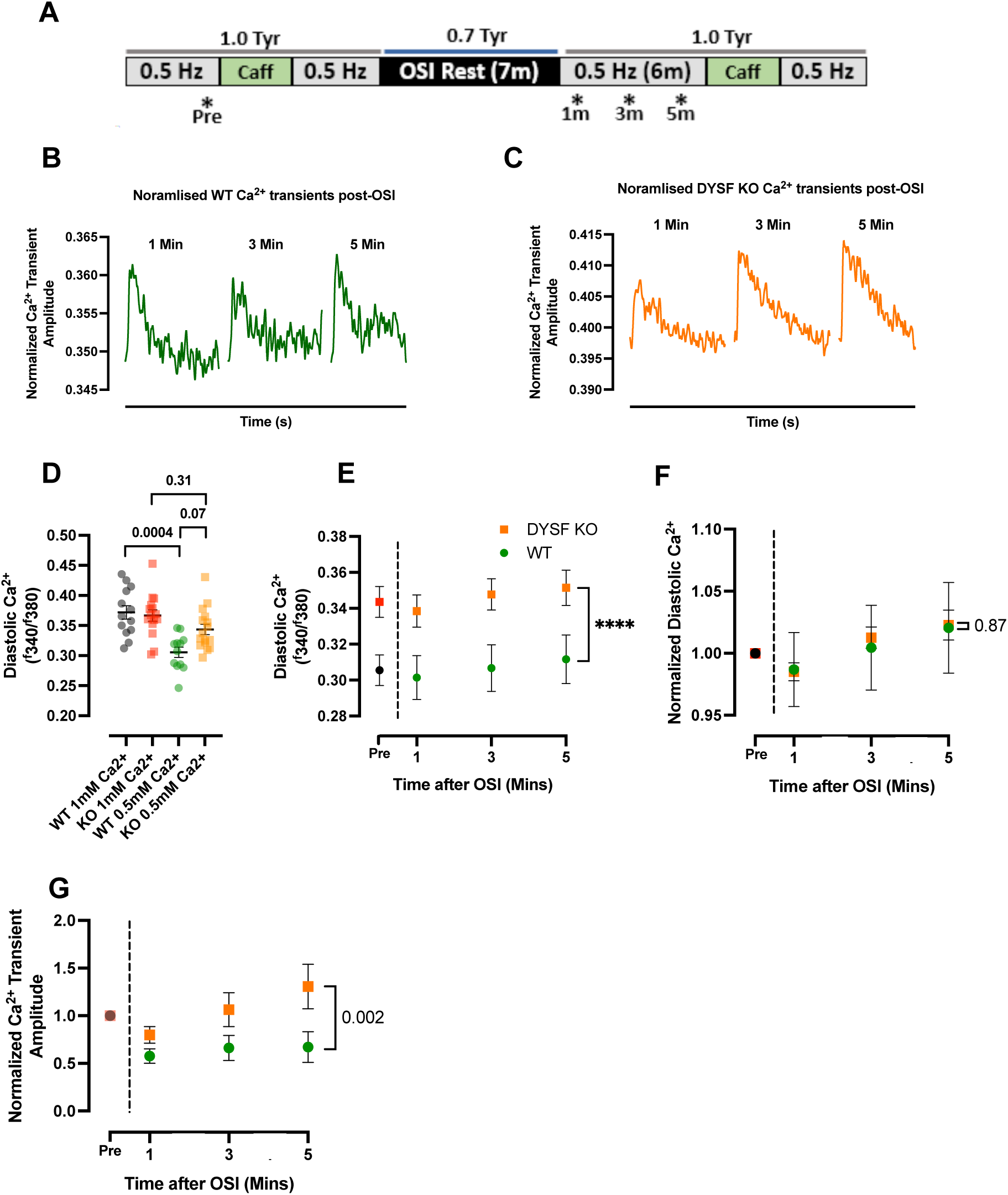
DYSF KO cardiac myocytes gain Ca^2+^ after osmotic shock injury and respond inappropriately to low extracellular Ca^2+^. **A,** A timeline demonstrating the *in vitro* OSI protocol. *’s indicate time point for data collection in **E**, **F** and **G**. B & C, representative average Ca^2+^ transients at 1, 3 and 5 minutes post-OSI in WT and DYSF KO cells, respectively. **D**, comparison of diastolic Ca^2+^ at 1mM and 0.5mM extracellular Ca^2+^. *N* = 5-7 animals and *n* = 12-16 cells. **C**, diastolic Ca^2+^ Pre-OSI and at 1-, 3- and 5-minutes after OSI. **E**, diastolic Ca^2+^ pre-OSI and at 1, 3 and 5 minutes post-OSI normalized to pre-OSI levels. **F**, Ca^2+^ transient amplitude at 1, 3 and 5 minutes post-OSI normalized to pre-OSI levels. **D-G** Data points represent genotype averages for each time point. WT – *N/n* = 5/12, KO – *N/n* = 7/16. A two-way ANOVA was used to determine statistical significance. Data is presented as mean ± SEM. **** - *P* = <0.0001.

We first investigated the effect of decreasing extracellular Ca^2+^ from 1- to 0.5mM. As expected, we observed a concurrent decrease in diastolic Ca^2+^ in WT cardiac myocytes. However, diastolic Ca^2+^ was unaltered in DYSF KO cells despite the halving of extracellular Ca^2+^, which lead to a trend towards increased diastolic Ca^2+^ in DYSF KO compared to WT cells in 0.5 mM Ca^2+^ (*p=0.*07; **Fig. 6D**). This indicates that DYSF KO cells are unable to adapt to changes in extracellular Ca^2+^. Throughout the immediate post-OSI period (5 mins) diastolic Ca^2+^ was elevated in DYSF KO cells relative to WT cells (**Fig. 6E**). However, relative to pre-OSI levels, we found that OSI brought about comparable changes in diastolic Ca^2+^ in both WT and DYSF KO cells during this time (**Fig. 6F**). On the other hand, while Ca^2+^ transient amplitude remained stable after OSI in WT cells, a progressive increase in Ca^2+^ transient amplitude was observed in DYSF KO cells, indicative of dysregulated Ca^2+^ handling in DYSF KO cells post-OSI (**Fig. 6G**). The lack of temporal changes in diastolic Ca^2+^ between genotypes (**Fig. 6F**) suggests that a difference in diastolic Ca^2+^ is not responsible for the progressive increase in Ca^2+^ transient amplitude in DYSF KO cells after OSI. To better understand this, we next investigated post-OSI Ca^2+^ handling characteristics in more detail.

### Dysregulation of intracellular Ca^2+^ handling leads to accumulation of Ca^2+^ in DYSF KO cells after OSI

During these low external Ca^2+^ experiments it was evident that the rapid application of 10mM caffeine used to measure SR Ca^2+^ content enhanced the following stimulated Ca^2+^ transients. Therefore, we compared the characteristics of Ca^2+^ transients at 0.5 Hz stimulation after cells had reached steady state following caffeine application. A diagram of the OSI protocol is shown in **Figure 7A** where ‘^’ indicates the pre- and post-OSI time points used to measure Ca^2+^ handling characteristics in **Figures 7B-Q**. Overall, OSI caused limited changes to the Ca^2+^ handling capabilities of WT cells (**Fig.7B-I**). Specifically, we observed a non-significant trend towards a decrease in Ca^2+^ transient amplitude after OSI in WT cells (**Fig. 7C**) with no change in systolic Ca^2+^ (**Fig. 7D**) diastolic Ca^2+^ (**Fig. 7E**) or the amplitude of the caffeine transient (indicative of SR Ca^2+^ content; (**Fig. 7F**). In the WT, OSI did produce a slowing of the rate of decay of the Ca^2+^ transient (*k*_sys_; **Fig. 7G**), which was due to decreased SERCA function (*k*_sr_; **Fig. 6H**) rather than a SERCA independent mechanism (*k*_caff_; **Fig. 7I**).

**Figure 7.**
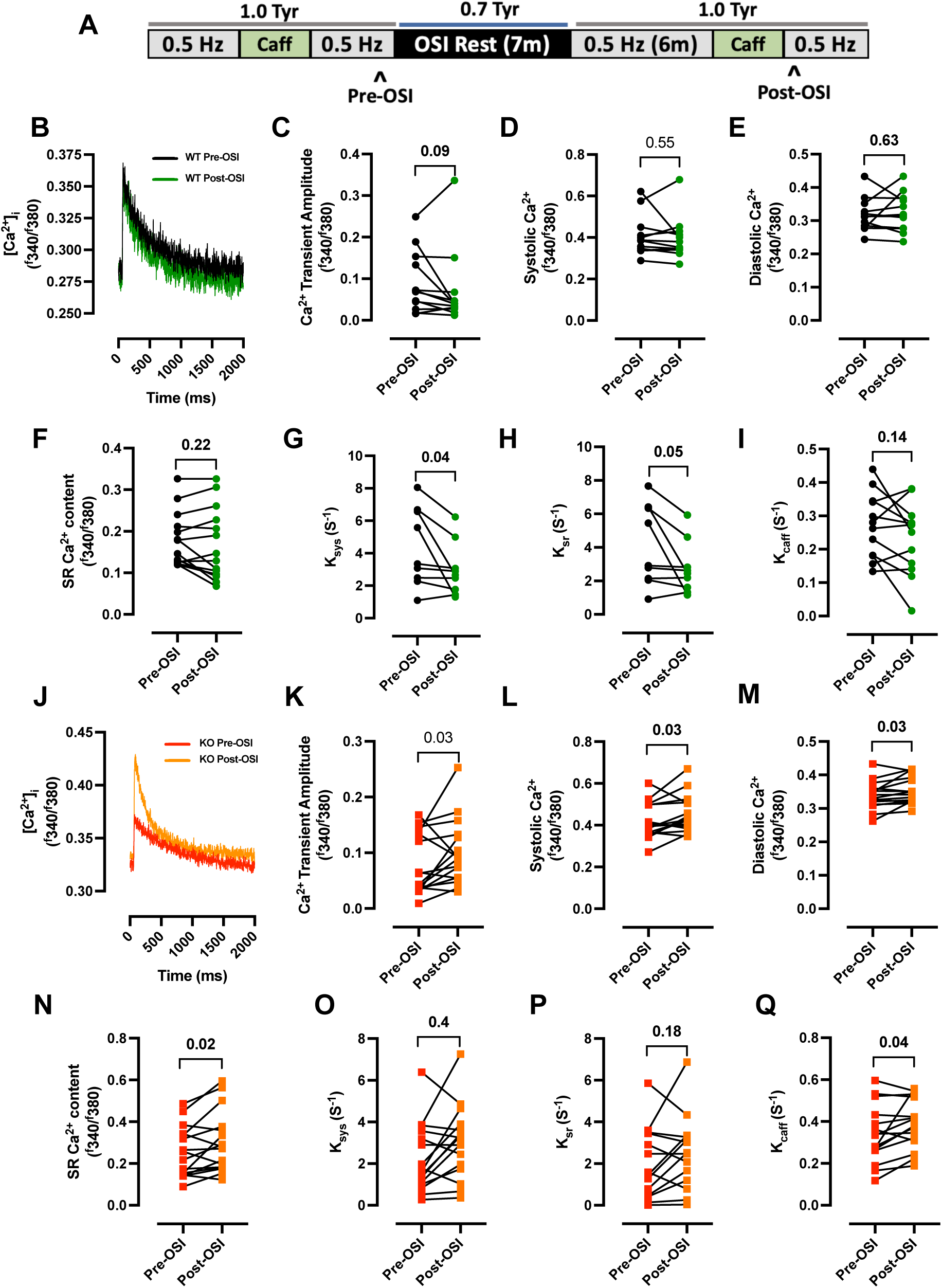
*In vitro* osmotic shock injury causes dysregulated Ca^2+^ handling in DYSF KO cardiac myocytes. **A,** A timeline demonstrating the *in vitro* OSI protocol. ^’s indicate time point for data collection in **B**-**Q**. **B**, Example Ca^2+^ transients in a WT cell pre- and post OSI. **C**, Ca^2+^ transient amplitude **D**, Systolic Ca^2+^ **E**, Diastolic Ca^2+^ **F**, SR Ca^2+^ content **G**, Rate constant of decay (RoD) of the Ca^2+^ transient (K_sys_) **H**, contribution of SERCA to Ca^2+^ transient RoD (K_sr_) **I**, RoD of the 10 mM caffeine transient (K_caff_) in WT cells pre-OSI vs. post-OSI. **J-Q**, same parameters as in **B-I** but showing DYSF KO cells pre-OSI vs. post-OSI. **C**, A Wilcoxon test and **D-I** & **K-Q** a paired non-parametric t-test were used to determine statistical significance. Data is presented as mean ± SEM.

In direct contrast to the WT cells, similarly to **Figure 6E**, we observed that in DYSF KO cells OSI increased Ca^2+^ transient amplitude (**Fig. 7K**) due to an increase in systolic Ca^2+^ (**Fig. 7L**) which predominated over an increase in diastolic Ca^2+^ (**Fig. 7M**). The SR Ca^2+^ content is a major determinant of the Ca^2+^ transient amplitude, and we found the amplitude of the caffeine evoked Ca^2+^ transient was increased upon OSI in DYSF KO cells **Fig. 7N**). In contrast to WT, in the DYSF KO cells OSI did not alter the rate of decay of the Ca^2+^ transient (*k*_sys_; **Fig. 7O**) or SERCA function (*k*_sr_; **Fig. 7P**) but the increase in Ca^2+^ transient amplitude was associated with an increase in *k*_caff_ suggesting an increase in NCX activity (**Fig. 7Q**). Therefore overall, our data suggests dysferlin protects cardiac cells from increased Ca^2+^ loading and *I*_NCX_ activity during acute stress due to OSI.

We next investigated if OSI exposed dysferlin sensitive differences in the synchronicity of the triggered Ca^2+^ release or the occurrence of Ca^2+^ sparks and waves using confocal line scan. The synchronicity of Ca^2+^ release was decreased in WT cells after OSI, consistent with decreased T-tubule density and increased fragmentation and we observed a similar non-significant trend in the DYSF KO cells (**Supplementary Fig. 3**), but the frequency of Ca^2+^ sparks or spontaneous global Ca^2+^ release events was not altered (**Supplementary Fig. 3**). Therefore, OSI-induced dysregulation of Ca^2+^ handling in DYSF KO cells does not appear to be due to perturbed RyR Ca^2+^ release.

In summary, in DYSF KO cells versus WT, as well as driving a decrease in T-tubule density and connectivity (**Fig. 5**), OSI also caused an accumulation of Ca^2+^ within the cell likely due to pathological dysregulation upon OSI, rather than positive compensatory mechanisms observed in WT cells. Since OSI caused T-tubule fragmentation in both WT and KO cells, although to a greater extent in WT, this may be indicative of dysferlin protecting against the dysregulation of cardiac EC coupling in OSI via direct regulation of EC coupling proteins and not solely via the regulation of T-tubule network structure.

### Cardiac dysferlin abundance decreases with age and may contribute to age-dependent decline of the T-tubule network

Finally, cardiac ageing correlates with a decline in cardiac function and a degradation of the T-tubule network (Kong et al. 2018; Lyu et al. 2021). However, our understanding of why T-tubules are remodelled with age is poor and our understanding of whether dysferlin is important in this process is almost completely lacking.

We first sought to examine whether dysferlin expression changes during ageing. As such, we observed a natural age-related decline by approximately 55% in cardiac dysferlin abundance at 10-months-of-age compared to 3-5 months of age (**Fig. 8A & Ai**). Since we have shown that dysferlin KO decreases density and increases fragmentation (**Fig. 1**), we set out to investigate if ageing exerts effects on cardiac T-tubules via the loss of dysferlin.

**Figure 8.**
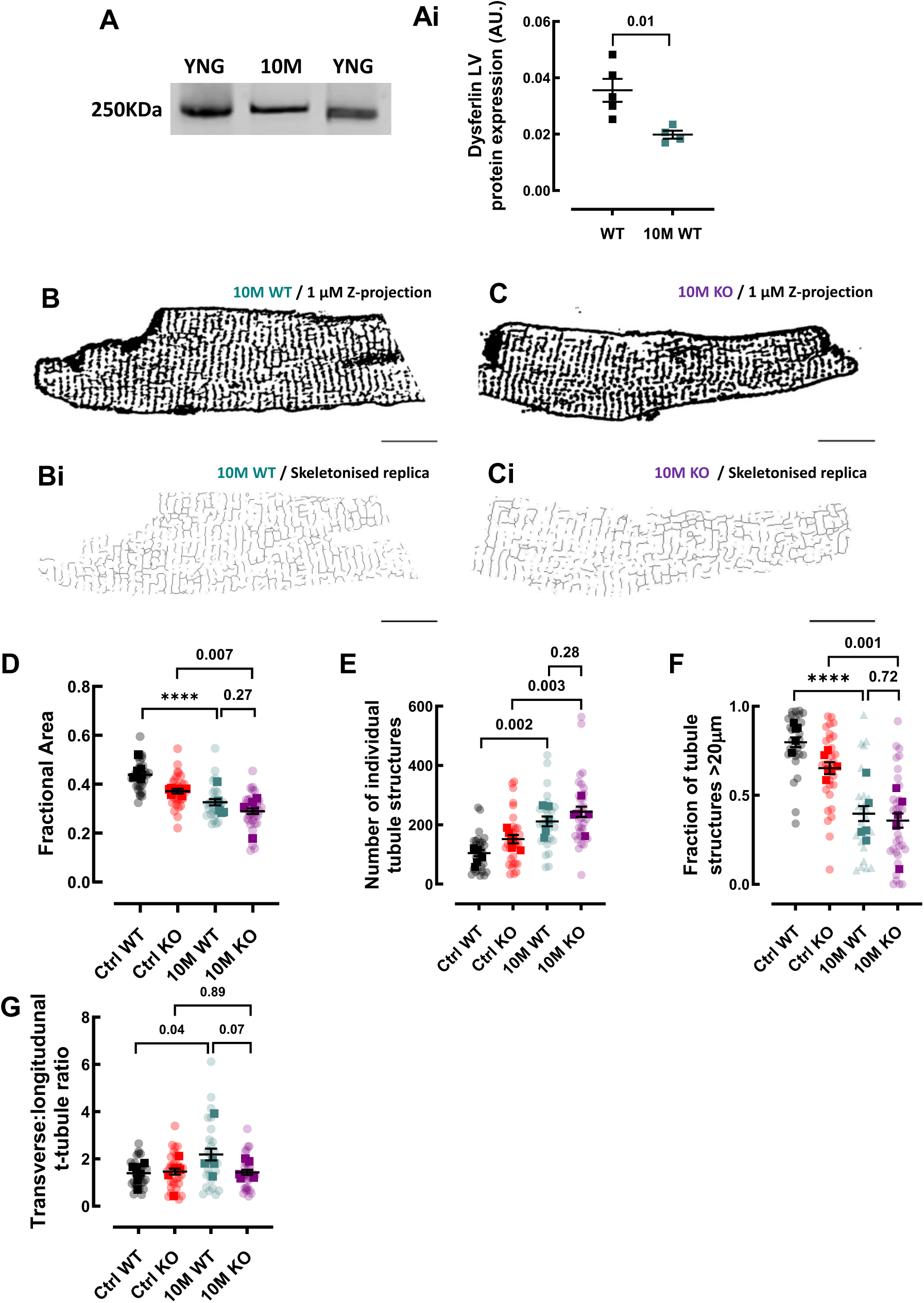
A natural reduction in cardiac dysferlin expression correlates with age-related T-tubule Decline. **A & Ai** Representative Western blot and left ventricular dysferlin protein expression in young WT and 10-month-old (10M) WT mice. **B & C**, Representative composite confocal images of the T-tubule network stained with Di-4-ANEPPS in isolated WT and DYSF KO mouse ventricular myocytes from 10-month-old (10M) mice. **Bi & Ci**, Skeletonised replicas of the T-tubule networks shown in **B & C**. **B-Ci**, Scale bars = 15 µm. **D**, T-tubule fractional area. **E**, Number of individual tubule structures. **F**, Fraction of tubule structures over 20 µm in length. **G**, Transverse/longitudinal tubule ratio. **A**, *N* = 4-5. An unpaired two-tailed t-test was used to determine statistical significance. **D - G**, *N* = 5-6 mice (opaque data points), *n* = 25-32 (translucent data points). Linear mixed model analysis was used to determine statistical significance. Data is presented as mean ± SEM. **** - *P* = <0.0001.

T-tubule networks in 10-month-old WT and DYSF KO cardiac myocytes were examined using confocal z-stacks. Representative images of 10-month-old WT and DYSF KO T-tubule networks are shown in **Fig. 8B & C**. Overall, compared to dysferlin KO in young mice, ageing produced similar, but more pronounced effects on the T-tubule network. Specifically, ageing decreased T-tubule density (**Fig. 8D**) and increased fragmentation of T-tubules (as measured by increased number of T-tubules and decreased large T-tubule structures; **Fig. 8E & F**) such that there was no difference between aged WT and dysferlin KO T-tubule networks (**Fig. 8E & F**). Interestingly, there was an increase in the ratio of transverse:longitudinal tubules with age in WT but not in DYSF KO cells, which may represent a compensatory mechanism in the WT, which is lacking in DYSF KO cells (**Fig. 8G**). Taken together, these findings suggest that a decrease in dysferlin abundance plays an important role in age-associated T-tubule remodelling.

## Discussion

In this study we have shown that dysferlin regulates the structure and function of the mouse cardiac EC coupling machinery under baseline conditions, in response to acute stress *in vitro* and during normal ageing, as well as increasing the arrhythmogenic burden in *ex vivo* hearts. Specifically, we found that under baseline conditions dysferlin loss, (i) decreased T-tubule density and connectivity, (ii) narrowed the dyadic cleft and (iii) decreased the Ca^2+^ transient amplitude and the rate of removal of cytosolic Ca^2+^. In addition, under conditions of stress dysferlin loss exacerbated cardiac responses by, (iv) increasing arrhythmias in the presence of isoprenaline, (v) increasing the susceptibility of T-tubules to OSI-induced fragmentation and (vi) caused OSI induced dysregulation of Ca^2+^ handling which did not occur in WT cells. Finally, we show (vii) a natural decline in cardiac dysferlin abundance is important for T-tubule degradation during normal ageing.

### Dysferlin is important for normal T-tubule density and connectivity

At present, little is known about cardiac T-tubule dynamics. However, we have shown that dysferlin loss decreases t-tubule density, resulting in a more fragmented and less branched T-tubule network in cardiac myocytes from young, otherwise healthy mice. This suggests dysferlin modulates physiological T-tubule turnover in the heart with the decrease in T-tubule connectivity upon dysferlin loss likely being detrimental. Maladaptive T-tubule remodelling has also been described in DYSF KO cardiac myocytes from 12 to 14month-old mice, however it is difficult to compare our findings from young animals with those observed using a moderately aged model (Hofhuis et al. 2020) since we show ageing itself decreases dysferlin.

A key player in T-tubule turnover is bridging integrator 1 (BIN-1), which is thought to drive the invagination of T-tubules from the sarcolemma in striated muscle cells (Caldwell et al. 2014; Lee et al. 2002; Perdreau-Dahl et al. 2023). To date dysferlin is the only other protein known to promote *de novo* tubule growth albeit in a non-cardiac cell line (Hofhuis et al. 2017). When dysferlin and BIN-1 are co-expressed *in vitro*, they act synergistically during tubule formation, where the authors suggest that dysferlin fine tunes t-tubule structure (Hofhuis et al., 2017). Our findings support this hypothesis as a relatively well-developed T-tubule network is still present in cardiac myocytes that lack dysferlin.

In addition to gross morphological changes in the DYSF KO T-tubule network, we also found significant ultrastructural disorder using transmission electron micrographs. The nanoarchitecture of the T-tubule membrane, such as the arrangement of membrane folds, has been shown to be important for the tight regulation of ionic flux during EC coupling (Hong et al. 2014). This suggests, therefore, that malformation within some of the DYSF KO T-tubules may partially contribute to the mishandling of Ca^2+^ we have observed in DYSF KO cardiac myocytes.

Although a topic of debate, dysferlin is thought to facilitate vesicle trafficking and fusion in muscle cells (For review see Quinn et al., 2024). Other members of the ferlin family are also known to mediate vesicle trafficking in non-muscle tissues (Lek et al. 2012). Vesicle recruitment likely facilitates T-tubule maturation in skeletal muscle (Lee et al. 2002) and a recent report suggests dysferlin-mediated vesicle trafficking drives tubule proliferation in the heart, which may underpin the onset of cardiac hypertrophy in response to experimental chronic pressure overload (Paulke et al. 2024). We hypothesize, therefore, that reduced dysferlin-mediated vesicle trafficking may contribute to maladaptive remodelling of the T-tubule network in DYSF KO cells. However, further investigation is required to confirm the contribution of dysferlin-containing vesicles to T-tubule turnover in striated muscle.

### Dysferlin is important in maintaining dyadic structure

Dysferlin is known to localise to the mouse T-tubule membrane (Hofhuis et al. 2020). More recent nanometric scale spatial analysis also localised dysferlin to the human cardiac dyad and suggested that JPH2, a key regulator of the dyad (Lehnart and Wehrens 2022), may actively recruit dysferlin to the dyadic cleft (Paulke et al. 2024). In skeletal muscle, dysferlin inserts into the T-tubule membrane via its C-terminal transmembrane domain (Kerr et al. 2013) and therefore, it is likely that dysferlin anchors to the SR membrane in the same way, although this has not yet been determined. Dysferlin likely exists in physical macromolecular complexes with the RyR2, which itself is embedded in the SR membrane (Paulke et al. 2024). These findings suggest that dysferlin could span the cardiac dyadic cleft or at least integrate into both the T-tubule and SR membranes.

To the best of our knowledge, we show here that dysferlin is the only protein known to date to modulate the width of the cardiac dyadic cleft. It is established that JPH2 is essential for dyad formation as it anchors the T-tubule and SR membranes in close opposition such that JPH2-KO in cardiac myocytes leads to a reduction in the formation of junctional membrane complexes (van Oort et al. 2011) but not the width of the complexes formed. There is a distinct absence of literature regarding how dyadic width is controlled and therefore, any suggestion as to how dysferlin may regulate the width of the dyadic cleft is speculative. Dysferlin has, however, been shown to have a minor axis width of ∼102 angstroms or 10nm (Huang et al. 2024), which would occupy much of the dyadic cleft width of 12-15nm. Thus, the absence of dysferlin may cause the T-tubule and jSR membranes to exist in closer proximity to each other thus narrowing the dyad. Whether the observed narrowing of the dyadic cleft affects Ca^2+^ release is unknown. Were this structural change to occur alone it would not be expected to produce a sustained change in systolic Ca^2+^ (Eisner et al., 1998). However, in the setting of other changes to the Ca^2+^ handling process we observe at baseline (decreased Ca^2+^ transient amplitude and fractional SR Ca^2+^ release) it is tempting to speculate that dyad narrowing could be compensatory. Whether dyadic narrowing could be detrimental and promote aberrant Ca^2+^ release and arrhythmias during conditions of stress e.g. where SR Ca^2+^ content is increased upon OSI, is unknown. The functional consequences of dyadic narrowing on Ca^2+^ release in DYSF KO cardiac myocytes is therefore uncertain warranting further investigation whilst also considering other potential functional modifications to key EC coupling proteins (Mattiazzi et al. 2006; Niggli et al. 2013).

### Dysferlin is important for setting the normal Ca^2+^ transient amplitude and rate of decay

In conjunction with decreased T-tubule density and connectivity in DYSF KO cardiac myocytes, the amplitude and rate of decay of the systolic Ca^2+^ transient was decreased. Dysferlin loss did not alter the amplitude of the caffeine-induced Ca^2+^ transient, suggesting no change in the SR Ca^2+^ content. Our data therefore points to the decreased fractional Ca^2+^ release from the SR in DYSF KO cells underpinning the decreased Ca^2+^ transient amplitude at baseline. An important regulator of both fractional SR Ca^2+^ release and Ca^2+^ transient amplitude is the L-type Ca^2+^ current (*I*_Ca-L;_ (Fearnley et al. 2011). LTCCs are primarily located on T-tubules where T-tubule loss is associated with decreased *I*_Ca-L_ (Clarke et al. 2015; Dibb et al. 2009; Kamp and He 2002). Hofhuis et al., reported a decrease in *I*_Ca-L_ in DYSF KO cardiac myocytes compared to WT controls in 14-month-old mice and in skeletal muscle dysferlin modulates LTCC function (Kerr et al. 2013; Lee et al. 2002; Lukyanenko et al. 2017; Lukyanenko et al. 2022). Quantification of *I*_Ca-L_ was outside the scope of our study but if a similar decrease in *I*_Ca-L_ occurs in DYSF KO cells at 3-5-months (where t-tubules are lost) this could explain the decrease in fractional SR Ca^2+^ release and the decreased Ca^2+^ transient amplitude we observe. In contrast to our findings in young mice, Hofhuis *et al*. reported an increase in fractional SR Ca^2+^ release in DYSF KO cells and an unchanged Ca^2+^ transient amplitude. The mice used in their study were moderately aged and therefore discrepancies with our own work using young animals may reflect the natural age associated loss of dysferlin we report here.

In addition, we show dysferlin loss slows the decay of the Ca^2+^ transient in agreement with findings by Wei et al., 2015 but in contrast to studies in moderately aged animals where no change was detected (Hofhuis et al. 2020). Our study shows the slowed decay of the Ca^2+^ transient upon dysferlin loss was due to decreased SERCA function as well as a SERCA independent pathway. Decreased SERCA function is well known to occur in numerous pathological conditions where it is associated with the negative force frequency relation seen in human heart failure and can also drive cardiac alternans, a precursor of arrhythmias (Hasenfuss et al. 1994). As such increasing SERCA function is a promising therapeutic target (for review see (Korpela et al. 2021)).

### Dysferlin regulates normal heart rhythm

Our findings are the first to show that dysferlin loss increases the susceptibility of the heart to sustained cardiac arrhythmias. Interestingly, the increased susceptibility of DYSF KO hearts to arrhythmias was unmasked upon stressing the heart with isoprenaline but not during baseline conditions, suggesting an inability of DYSF KO hearts to respond appropriately.

Precisely how the lack of dysferlin increases arrhythmia susceptibility remains unknown however, the generation of arrhythmias requires both a triggering mechanism and a substrate upon which an arrhythmia can propagate (Keren et al. 1984). Since arrhythmias were increased in isoprenaline by an increase in sustained events rather than any change in premature ventricular complexes (single, double or triplet) our data suggests dysferlin loss promotes the conversion of premature events into sustained arrhythmias. This could occur by structural remodelling, commonly an increase in fibrosis (de Jong et al. 2011) however, low and comparable levels of myocardial fibrosis have been observed in young dysferlin-deficient mice compared to WT controls (Chase et al. 2009) making this mechanism unlikely to drive arrhythmias in this study.

On the other hand, disturbances in T-tubule structure and Ca^2+^ handling are also known to play a role in the maintenance of arrhythmias and therefore we speculate that they could play a role in the DYSF KO mouse. T-tubule loss and disruption occurs in DYSF-KO hearts but particularly in these hearts during stress induced by OSI; such remodelling is known to cause heterogeneous Ca^2+^ release which can increase the occurrence of arrhythmias (Biesmans et al. 2011; Louch et al. 2004; Singh et al. 2017) for review see (Dibb et al. 2022). A Ca^2+^ dependent arrhythmia mechanism linked to stress (isoprenaline) is consistent with the exacerbated dysregulation of Ca^2+^ handling during stress (OSI) seen in the DYSF-KO mouse. While WT hearts respond to OSI by decreasing SERCA function, in DYSF-KO hearts SERCA was unchanged by OSI and hearts suffered from an increase in intracellular Ca^2+^. This was characterised by an increase in SR Ca^2+^ content (caffeine transient amplitude) and the decay of the caffeine evoked Ca^2+^ transient, indicative of an increase in NCX function. Both increased SR Ca^2+^ content and NCX function are hallmark features of arrhythmias (Diaz et al. 1996; Sipido et al. 2000; Venetucci et al. 2008) and therefore may be involved in the arrhythmia susceptibility of the DYSF-KO heart.

### Dysferlin protects T-tubule structure during acute stress by Osmotic Shock Injury

This study is the first to show that dysferlin protects cardiac T-tubules and Ca^2+^ handling from acute stress induced by OSI and is consistent with dysferlin’s reputation in skeletal muscle as a membrane repair and protective protein (For Review see Quinn et al., 2024). OSI was chosen due to its pathophysiological relevance to conditions like ischemia or the application of cardioplegic solutions (Handy et al. 1996; Moench et al. 2013; Vandenberg et al. 1996). OSI damages t-tubules due to a rapid sealing of the tubule lumen upon resolution of hypo-osmotic shock (when cells shrink back to a normo-osmotic volume); it is thought that the shrinking causes the internalization of ‘membrane blebs’ into the T-tubule lumen due to the disconnection of T-tubule membrane from the surrounding cytoskeleton (Moench et al. 2013; Uchida et al. 2020). The increased susceptibility of DYSF-KO t-tubules to OSI induced fragmentation could therefore arise due to decreased connectivity between dysferlin-KO T-tubules and the intracellular architecture (Uchida et al., 2020).

Our novel findings in cardiac myocytes are consistent with the skeletal muscle literature. In skeletal muscle OSI traps membrane impermeable dyes within the dysferlin-null skeletal muscle T-tubule system (Kerr et al. 2013). This is consistent with OSI-induced T-tubule sealing, which we suggest is comparable to the proposed OSI-mediated T-tubule sealing mechanism in cardiac myocytes. Interestingly, either the removal of extracellular Ca^2+^ or pharmacological inhibition of the LTCC with diltiazem prevented the OSI-mediated T-tubule sealing in dysferlin-null skeletal muscle fibres (Kerr et al. 2013). This observation is particularly important considering our own findings that show DYSF KO cardiac myocytes accumulate Ca^2+^ after OSI. Thus, we hypothesize that dysferlin protects the cardiac and skeletal muscle T-tubule system from acute stress via a shared mechanism, which is dependent on preventing the pathological influx of extracellular Ca^2+^.

### Dysferlin protects cardiac myocytes from Ca^2+^ accumulation following acute stress by Osmotic Shock Injury

Our data shows for the first time that dysferlin regulates intracellular Ca^2+^ handling in cardiac myocytes in response to acute stress. Except for a change in SERCA function, OSI did not alter Ca^2+^ handling in WT cardiac myocytes, however there was pronounced OSI-induced dysregulation of intracellular Ca^2+^ handling in DYSF KO cells. Since OSI did alter WT T-tubules (although to a lesser extent than in the KO) with little effect on Ca^2+^ handling the pronounced effect on Ca^2+^ handling in DYSF-KO cells may point to a role for a t-tubule independent as well as t-tubule dependent factors.

Our experiments revealed that unlike WT cells, DYSF KO cells were unable to reduce diastolic Ca^2+^ when extracellular Ca^2+^ was decreased from 1 to 0.5mM. Since the purpose of decreasing extracellular Ca^2+^ experimentally was to limit cell death upon OSI this may have contributed to the dysregulated Ca^2+^ handling seen in DYSF-KO post-OSI. The reasons for this apparent inability to adapt to changing Ca^2+^ levels are unclear but may be linked to the decrease in SERCA dependent and independent Ca^2+^ efflux in DYSF KO cells at baseline such that decreased Ca^2+^ efflux pathways are unable to manage simple Ca^2+^ dependent manoeuvrers. This phenomenon resulted in increased diastolic Ca^2+^ in DYSF KO cells throughout the OSI protocol. However, during the immediate 5-minute post-OSI period, on top of the increased diastolic Ca^2+^ which progressively increased with time, Ca^2+^ transient amplitude also increased progressively in DYSF KO cells compared to WT controls due to increased systolic Ca^2+^ levels. This suggests DYSF KO cells gain Ca^2+^ in response to OSI whereas WT cells effectively regulate Ca^2+^ levels. Similarly, SR Ca^2+^ content was increased by OSI in DYSF KO but not WT cells, which supports the concept of DYSF KO cells (but not WT cells) accumulating Ca^2+^ after OSI. Since we saw no evidence of increased SR Ca^2+^ leak in DYSF KO cells after OSI, the increase in diastolic Ca^2+^ after OSI is likely due to an influx of Ca^2+^ from the extracellular environment. Increased influx from the extracellular environment would also explain the increase in SR Ca^2+^ content, which occurred in the absence of any OSI-dependent change in SERCA function.

In contrast to our findings in DYSF KO cardiac myocytes, *in vitro* OSI injury caused a reduction in Ca^2+^ transient amplitude in dysferlin-null skeletal muscle fibres compared to WT controls (Kerr et al. 2013; Lukyanenko et al. 2017). This is due to the increased SR Ca^2+^ leak observed in dysferlin-null skeletal muscle fibres, which reduces SR Ca^2+^ content and thus the amount of Ca^2+^ released upon stimulation. A lack of RyR2 leak after OSI in DYSF KO cardiac myocytes allows SR Ca^2+^ content and Ca^2+^ transient amplitude to increase as a consequence of Ca^2+^ accumulation within the cell. Although removal of extracellular Ca^2+^ protects dysferlin-null skeletal muscle T-tubules from OSI-mediated fragmentation, removal of extracellular Ca^2+^ does not rescue OSI-induced Ca^2+^ mishandling (Lukyanenko et al. 2017). Therefore, the authors suggest that the destabilization of the biomechanical coupling between the LTCC and RyR1 underpins the OSI-mediated Ca^2+^ handling pathology in dysferlin-null muscle fibres. They also suggest that treatment with either LTCC or RyR1 blockers both protect against OSI-induced Ca^2+^ pathology due to a reduction in RyR1 Ca^2+^ leak (Lukyanenko et al. 2017).

It is clear that dysferlin loss leads to disparate effects on Ca^2+^ handling in response to acute stress in the form of osmotic shock injury in cardiac vs. skeletal muscle. These findings are of significant clinical relevance when considering why dysferlinopathy patients generally present with more severe skeletal muscle dysfunction compared to cardiac complications. Our data together with the current literature strongly suggests that dysferlin protects against the dysregulation of Ca^2+^ handling in response to stress independently of its role in T-tubule dynamics in both striated muscle types.

### An age-related decrease in dysferlin is important for T-tubule disruption in ageing

Finally, very little is known regarding the natural ageing process of cardiac T-tubules although independent reports show that T-tubule density declines with age (Kong et al. 2018; Lyu et al. 2021). We have shown for the first time that dysferlin is involved in the age-related t-tubule loss. Dysferlin KO altered t-tubule density and structure in young animals. However, the age associated decrease in WT cardiac dysferlin levels, by ∼55% in WT animals by 10-months-of-age, resulted in similar T-tubule networks in 10-month WT and DYSF KO mice suggesting the dysferlin loss in WT drives this age effect. Thus, we believe our data fosters the hypothesis that a natural reduction in cardiac dysferlin abundance is important for the age-related T-tubule decline. Hofhuis et al., compared the T-tubule networks of WT and DYSF KO mouse cardiac myocytes at a single 14-month-old time point and in contrast to our findings observed extensive structural deficiencies within DYSF KO T-tubule network. It is unclear whether WT dysferlin expression remains low during this time or whether other factors cause increased T-tubule remodelling in the DYSF KO between 10 and 14 months.

In our study, ageing to 10 months in WT mice resulted in an increase in transversally orientated tubules which did not occur in the DYSF-KO mouse. In young mice (3-5 months) genotype had no effect on T-tubule orientation in agreement with similar data at 8 months (Paulke et al. 2024) suggesting the age associated orientation change occurs between 8 and 10 months in WT mice. While ageing wasn’t investigated, Hofhuis et al., reported more transversely oriented tubules in 14-month-old WT compared to DYSF KO cells. Taken together with our own work this is further supports an age associated orientation change which requires dysferlin. There is debate in the literature as to whether dysferlin supports the stability of transverse tubules or if it preferentially builds longitudinal networks (Hofhuis et al. 2020; Paulke et al. 2024). Our own work suggests that differences in age between studies may have contributed to discrepancies and the idea that dysferlin interacts preferentially with branches of different orientations and warrants further investigation.

## Summary

Here we have presented novel findings that show that dysferlin is a key regulator of cardiac structure and function. We show that dysferlin is important for T-tubule structure in healthy, young cardiac myocytes, in acute stress in response to hypo-osmotic stress and during ageing. We have also shown that dysferlin regulates the width of the cardiac dyad and that dysferlin is essential for healthy intracellular Ca^2+^ release under baseline conditions and in response to acute stress. Finally, we have shown for the first time that the loss of dysferlin renders the heart more susceptible to sustained ventricular arrythmias in response to experimental pacing. These data provide a basis for new innovative experiments to better understand the role of dysferlin in cardiac function in health and disease.

## Notes

### Competing Interest Statement

The authors have declared no competing interest.

### Funding

- British Heart Foundation: PG/22/10902

