## Supplementary Figures for "Dysferlin Regulates Cardiac T-tubule Structure and Excitation-contraction Coupling in Isolated Cardiac Myocytes at Rest and in Response to Acute Hypo-osmotic Stress and is Protective Against Arrhythmias in Langendorff-perfused Hearts"

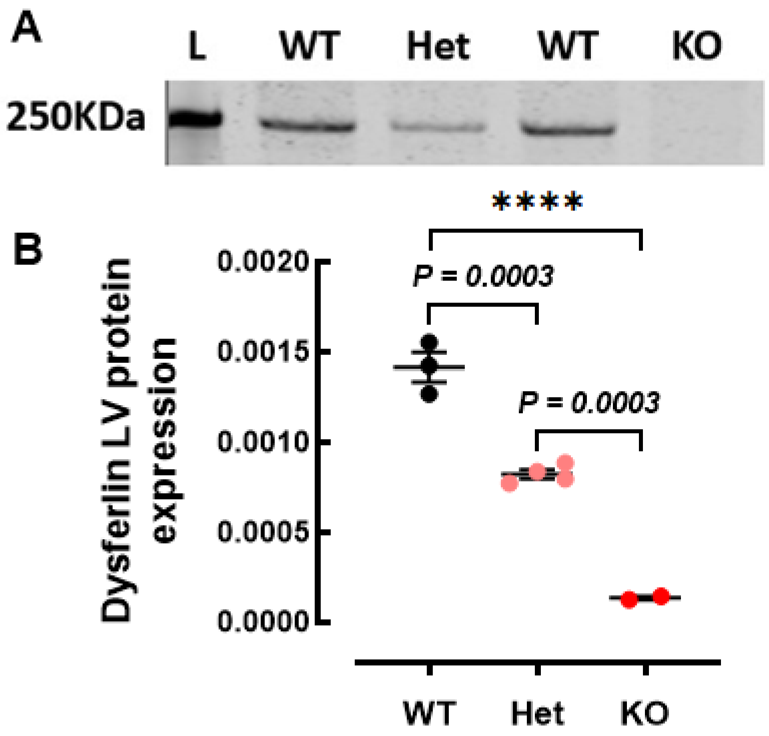


**Supplementary Figure 1. Confirmation of dysferlin loss in DYSF KO mouse model. A & B**, Representative Western blot stain and mean dysferlin protein expression in left ventricular myocardial tissue from wild type (WT), heterozygous mutant (Het) and DYSF KO mice. Three technical repeats were performed. *N* = 2-4. Statistical significance was determined using an ordinary one-way ANOVA with a post hoc Tukey's multiple comparisons test. **** = $P < 0.0001$.

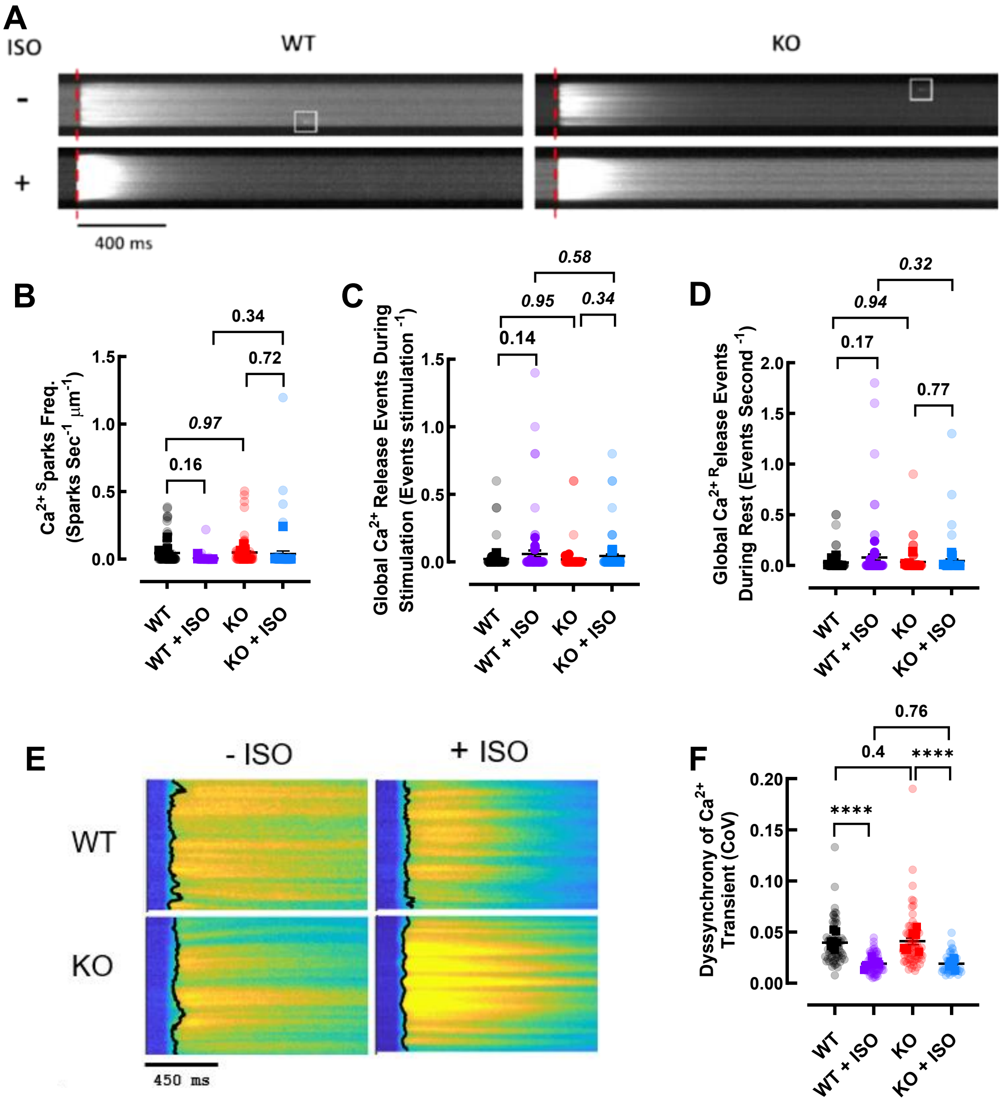


**Supplementary Figure 2. $Ca^{2+}$ Spark Frequency, Premature Global $Ca^{2+}$ Release Events and Dyssynchrony Index of $Ca^{2+}$ Release at Baseline**. **A**, Representative confocal line scans showing a single $Ca^{2+}$ transient in WT and DYSF KO isolated mouse ventricular myocytes. Experiments were performed on cells loaded with a $Ca^{2+}$ sensitive dye (Fluo-5F) with or without the use of isoprenaline (ISO; 100nM) in 1mM $Ca^{2+}$. Examples of $Ca^{2+}$ sparks are indicated by white boxes. Scale bar = 400 ms. **B**, Average frequency of $Ca^{2+}$ sparks (Sparks $Sec^{-1}$ $\mu m^{-1}$) during a 10 second rest period after 5 stimulations at 0.5 Hz. **C**, Incidence of global $Ca^{2+}$ release events during 0.5Hz stimulation (Events Stimulation $^{-1}$). **D**, Incidence of global $Ca^{2+}$ release events during rest period (Events Second $^{-1}$) **E,** Example images of stimulated $Ca^{2+}$ transients used to measure dyssynchrony of $Ca^{2+}$ release. **F**, Dyssynchrony index (coefficient of variation / CoV) of $Ca^{2+}$ transient. *N* = 5-7 mice (opaque data points), *n* = 20-81 cells (translucent data points). Linear mixed model analysis was used to determine statistical significance. Data is presented as mean ± SEM. **** - *P* = <0.0001.

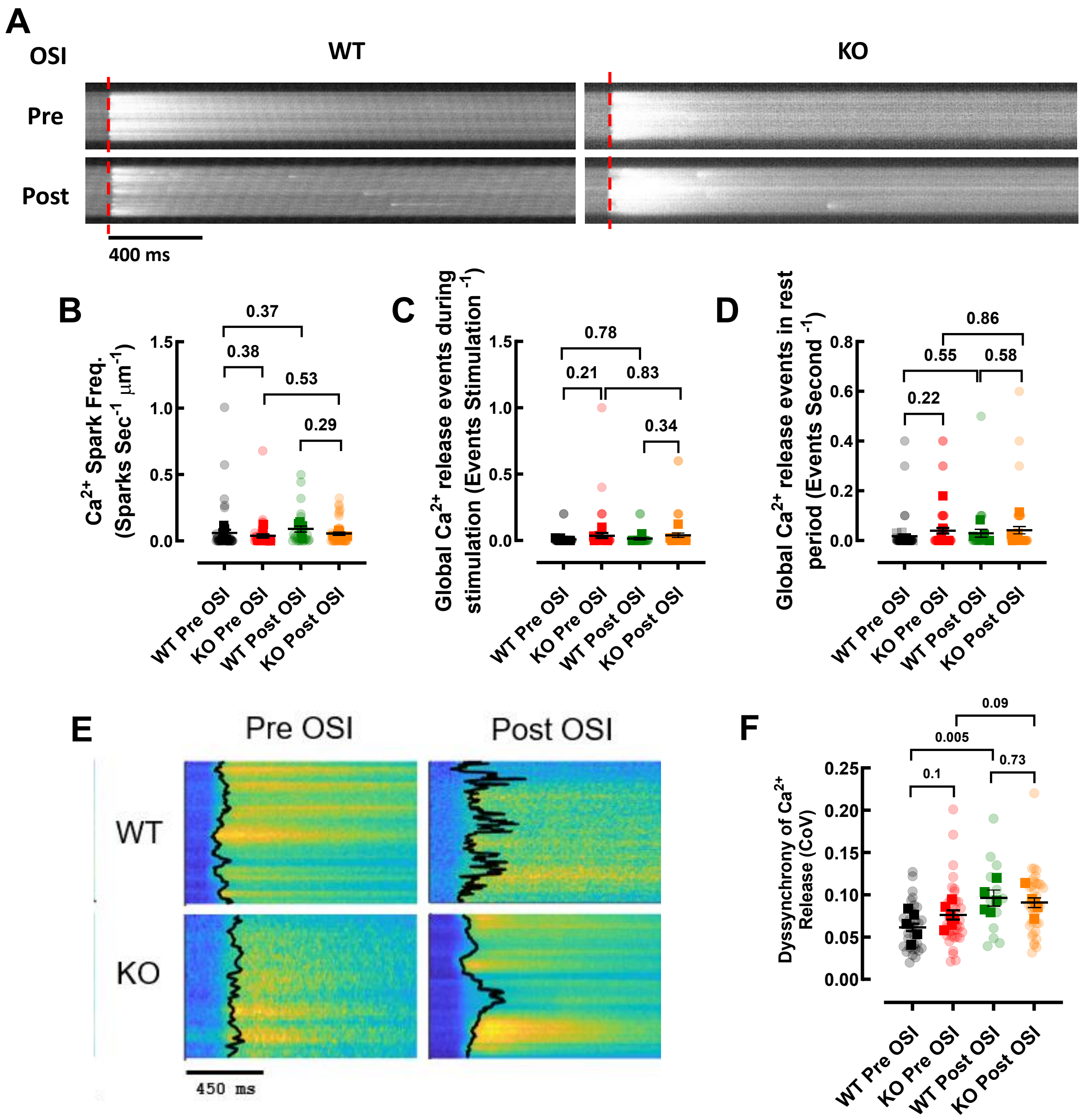


**Supplementary Figure 3. $Ca^{2+}$ Spark Frequency, Premature Global $Ca^{2+}$ Release Events and Dyssynchrony Index of $Ca^{2+}$ Release After OSI. A,** Representative confocal line scans showing a single $Ca^{2+}$ transient in WT and DYSF KO isolated mouse ventricular myocytes. Experiments were performed on cells loaded with a $Ca^{2+}$ sensitive dye (Fluo-5F) using 0.5mM $Ca^{2+}$ pre and post osmotic shock injury. Scale bar = 400 ms. **B**, Average frequency of $Ca^{2+}$ sparks (Sparks $Sec^{-1}$ $\mu m^{-1}$) during a 10 second rest period after 5 stimulations at 0.5 Hz. **C**, Incidence of global $Ca^{2+}$ release events during 0.5Hz stimulation (Events Stimulation $^{-1}$). **D**, Incidence of global $Ca^{2+}$ release events during rest period (Events Second $^{-1}$). **E**, Example images of stimulated $Ca^{2+}$ transients used to measure dyssynchrony of $Ca^{2+}$ release. **F**, Dyssynchrony index (coefficient of variation / CoV) of $Ca^{2+}$ transient. *N* = 5-6 mice (opaque data points), *n* = 12-49 cells (translucent [illegible]
